# Production the industrial levels of bioethanol from glycerol by engineered yeast “Bioethanol-4^th^ generation”

**DOI:** 10.1101/2020.06.04.132589

**Authors:** Sadat Mohamed Rezk Khattab, Takashi Watanabe

## Abstract

Besides the pledges for expanding uses of biofuels to sustain the humanosphere, abruptly massive needs emerged for sanitizers with turns COVID-19 to a pandemic. Therefore, ethanol is topping the social-demanding, although the three generations of production, from molasses/starch, lignocelluloses, and algae. Owing to the limited-availability of fermentable sugars from these resources, we addressed glycerol as a fourth bio-based carbon resource from biodiesel, soap, and fatty acid industries, which considers as a non-applicable source for bioethanol production. Here, we show the full strategy to generate efficient glycerol fermenting yeast by innovative rewriting the oxidation of cytosolic nicotinamide-adenine-dinucleotide (NADH) by O_2_-dependent dynamic shuttle while abolishing glycerol biosynthesis route. Besides, imposing a vigorous glycerol-oxidative pathway, the engineered strain demonstrated a breakthrough in conversion efficiency (up to 98%). Its capacity extending to produce up to 90g /l ethanol and > 2 g 1^-1^ h^-1^, which promoting the industrial view. Visionary metabolic engineering here provides horizons for further tremendous economic and health benefits with assuring for its enhancing for the other scenarios of biorefineries.

**Summary:** Efficiently fermenting glycerol in yeast was developed by comprehensive engineering the glycerol pathways and rewriting NADH pathways.

---

One of the challenges for sustaining the future humanosphere is producing adequate bio-based chemicals and fuels from renewable resources with the footprint toward reducing greenhouse gas emissions. The paradigm of using advanced sciences with metabolic engineering and biotechnologies for apt emerging needs of biofuels, materials, and chemicals has been envisioned and created on a commodity scale^1–3^. An abruptly massive needs in ethanol arose for medical uses as sanitizers, with turns COVID-19 to a pandemic; it had confirmed the efficiencies of 62-71% of ethanol for deactivating infection of the viruses’ attached to the hands and ward-off the infectious germs on persistent inanimate surfaces like metal, glass, and plastic^4^. Baker’s yeast (*Saccharomyces cerevisiae*), has several superior characteristics such as the ancient history with the safety of use, unicellular structure, short life cycle, distinguished powers of fermentation, robustness against inhibitors, stress-tolerance during different industrial levels of production, global infrastructures for production of bioethanol from starch and molasses, and the availability the toolboxes of genetic recombination. Besides, it is subjecting to the adaptive evolutions or even the hybridization, thence a Baker’s yeast had appointed as a top model platform of microbial cell factories for several biotechnological applications^5–7^. The first generation of bioethanol globally has successfully established with its uses for blending with gasoline as transportation biofuel. Owing to environmental, political, security, bio-economic issues, the demanding for bioethanol increases, although the resources for fermentation limited and the attempts are still enduring of overcoming the drawbacks of application of second and third generation of bioethanol from lignocellulosic biomass and the algae; basically, through evolving the maximum efficiencies in ethanol production during xylose fermentation with glucose or even coupled to acetic acid^8–12^.

In the last decade, glycerol producing industries, especially biodiesel, have expanded and accumulated substantial quantities of glycerol, which led to dropping its price^13^. Although the reductive merit in glycerol (C_3_H_8_O_3_) higher than other fermentable sugars^14^, glycerol is classifying as a non-fermentable carbon in the native *S. cerevisiae*^5^, besides, it is used poorly as feedstock, mainly through the glycerol 3-phosphate pathway, referred to as G3P pathway here, which composed of glycerol kinase (GUT1), and FAD-dependent-mitochondrial-glycerol-3-phosphate-dehydrogenase (GUT2)^15^. Conversely, yeast biosynthesizes glycerol for mitigating the osmotic stress and optimize the redox balance^16^, with subjection to the repression and transcriptional regulation of glucose through respiratory factors (RSF), and GUT1 and GUT2 genes^17–20^. The importance of glycerol as a carbon source, which could be utilized by yeast cells, has recognized. It promoted a study of the relationship between the molecular inheritance and the physiology of glycerol uptake and its metabolism. This study revealed a high interspecies diversity ranged from the good-glycerol grower to negative-glycerol grower in 52 of *S. cerevisiae* strains on a synthetic medium without supporting supplements and that the glycerol growth phenotype is a quantitative trait. It has confirmed that GUT1 is one of these genetic loci that sharing glycerol growth phenotype in one of these good-glycerol grower strains, a haploid segregant CBS 6412-13A^21^. Hereafter, two further superior alleles of cytoplasmic-ubiquitin protein-ligase-E3 (UBR2) and cytoplasmic-phosphorelay-intermediate osmosensor and regulator (SSK1) had found to link with GUT1 for the growing on the synthetic medium without supporting supplements^22^. These pivotal roles of UBR2 and GUT1 during glycerol assimilation by yeast had further confirmed by another study that re-sequenced the whole-genomes a glycerol-evolved strains^23, 24^ Although G3P-pathway has evidenced the main catabolic-pathway for glycerol catabolism in *S. cerevisiae*, its heterologous-replacing with DHA-pathway that combined glycerol facilitator (FPS) resulted in restores the similar growth of the parental strain. Furthermore, this replacement in a negative-glycerol grower strain bearing the swapped UBR2_CBS6412-13A_ allele had guided the growth rate to the highest specific growth rate ever reported on glycerol-synthetic medium^25^.

With an approach for the production of 1, 2-propanediol from glycerol, a significant amount of ethanol (18 g/l) had accumulated during the first day as a byproduct, particularly on the rich media. This study addressed metabolic engineering strategy combined heterologous-replaces of the G3P route by DHA-FPS pathway^25^ with a module for the production of 1, 2-propanediol, besides, the down-expression to the gene triosephosphate-isomerase gene (TPI1)^26^. Limiting oxygen availability in the shake flask cultures showed increasing the production of ethanol from glycerol (8.5 g ethanol / 51.5 g glycerol) to (15.7 g ethanol / 45 g glycerol) with production rate 0.1g l^-1^h^-1^ on synthetic medium in a recent study for facilitating understanding the future engineering of valuable products more reduced than ethanol^27^ using genetic modifications of heterologous-replaces of the G3P route by DHA-FPS pathway^25^. It is worth emphasizing glycerol has considered a non-fermentable carbon source in *S. cerevisiae*^5^; although, such attempts for fermenting it by *S. cerevisiae*. These experiments had initiated by overexpressed a native oxidative-glycerol pathway (DHA), includes glycerol dehydrogenase (GCY1) and dihydroxyacetone-kinase (DAK), beside overexpressed a glycerol uptake protein (GUP1) to produce 0.12g ethanol/ g glycerol with 0.025 g l^-1^h^-1^ of production rate^28^. Moreover, the methylotrophic yeast, *Ogataea polymorpha*, had tested for producing bioethanol from glycerol by overexpressing the genes involved either in the DHA or G3P pathways with integration with a gene of glycerol transporter FPS1 from *Pichia pastoris*. Furthermore, the recipient strain subjected to overexpress its genes of pyruvate decarboxylase (PDC1) and alcohol dehydrogenase (ADH1). Nonetheless, the overall ethanol produced was relatively low (10.7 g ethanol as a maximum accumulated product and 0.132g ethanol/ g glycerol) ^29^. Up to date, there is no native or genetically engineered strain promoting the industrial application of ethanol production from glycerol.

On the other hand, we developed a novel pretreatment method for biomass using glycerolysis with the catalysis of alum AlK(SO_4_)_2_, with additionally promoted by a microwave^30^. Hence, there emerged a need for evolving a model of yeast that can ferment glycerol efficiently after this glycerolysis for complete establishing our scenario by synergist current 4^th^ generation of bioethanol with its analog of the second or third generation, as well as either first generation. In this study, we report the details of how is the modeling of yeast cell to redirect the glycerol traffic to bioethanol production until the industrial levels even in the presence of glucose through the innovation of the forthcoming systematic metabolic engineering showed in (Fig 1): I) abolishing the inherent glycerol biosynthesis pathway by knocking-out NAD-dependent glycerol 3-phosphate dehydrogenase (GPD1) and retaining the second isoform GPD2 for requirements of glycerol 3-phosphate for lipid metabolism. II) Replacing cytosolic NADH-oxidation through the GPD1 shuttle by a more effective O_2_-dependent dynamic shuttle of water forming NADH-oxidase (NoxE) to renovate NAD^+^ for that integrated gene of glycerol dehydrogenase (GDH). III) Knocking out the first gene of the G3P pathway (GUT1). IV) Imposing a vigorous oxidative pathway via overexpressing two copies of both the heterologous-genes of glycerol dehydrogenase *Op*GDH, and the glycerol facilitator *Cu*FPS1, besides, the endogenous genes of TPI1, and DAK1 with one copy of DAK2.

**Fig.1.**
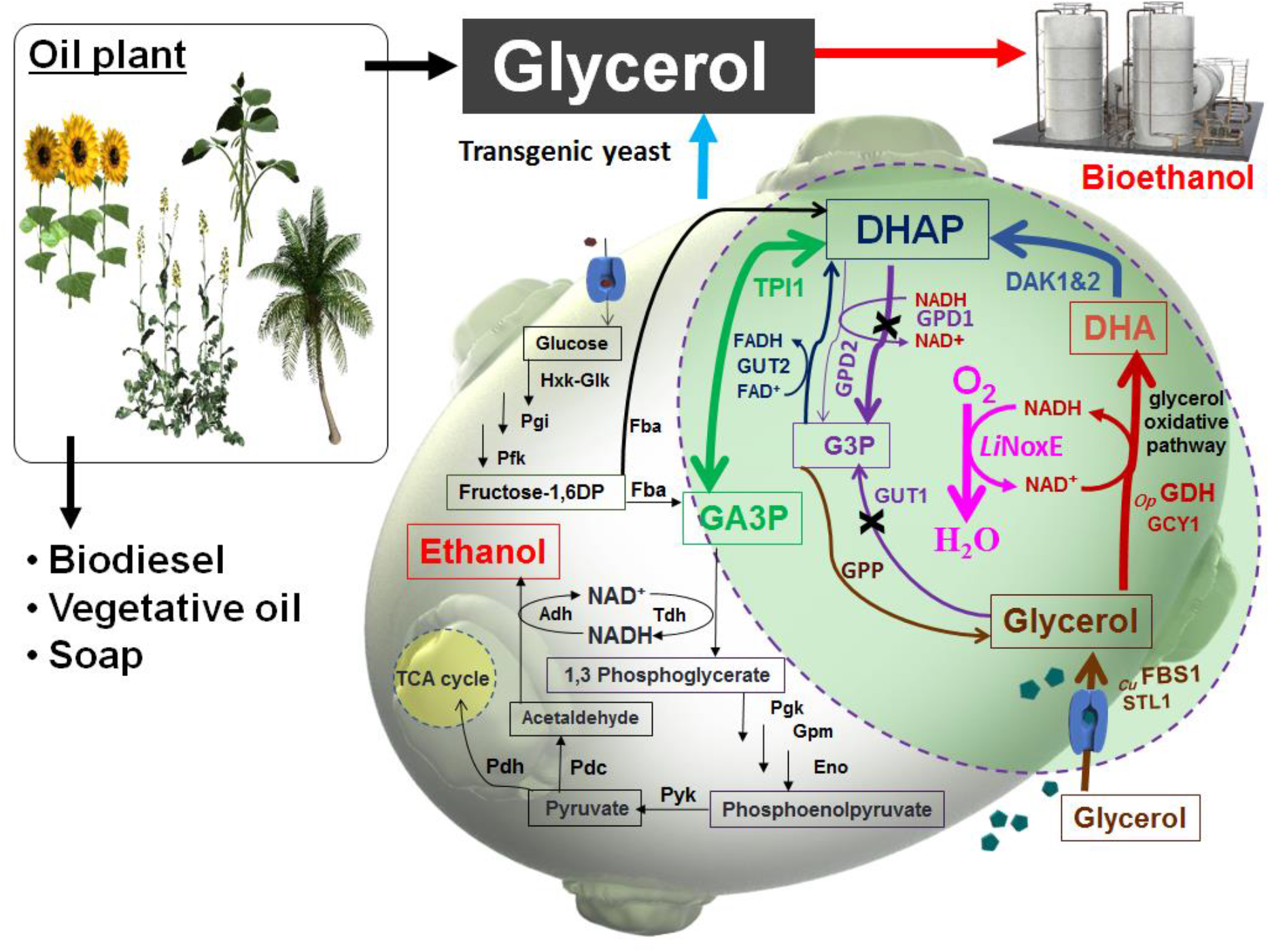
Schematic diagram showing the integrative scenario of biorefinery with a new generation of glycerol fermenting yeast, and redirection of glycerol influxes to ethanol production in *Saccharomyces cerevisiae via* retrofitted native glycerol anabolic and catabolic pathways with the robust oxidative route with renovation NAD^+^ cofactor by O_2_-dependent dynamic of water-forming NADH oxidase. During the pathway re-routing, glycerol-3-phosphate dehydrogenase (GPD1) and glycerol kinase (GUT1) were knocked out, bold arrows showing the overexpressed enzymes indigenous dihydroxyacetone kinase (*Sc*DAK1&2), triosephosphate isomerase (*Sc*TPI1), heterologous glycerol dehydrogenase from *Ogataea polymorpha* (*Op*GDH), glycerol facilitator from *Candida utilis* (*Cu*FPS1) and water-forming NADH oxidase from *Lactococcus lactis* subsp. *lactis* Il1403 (*Ll*NoxE).

## Results

### Effect of Systematic metabolic engineering

#### Step no. 1: vigorous glycerol dehydrogenase is an essential opener to initiate glycerol fermentation

Initial verification for overexpressing of glycerol dehydrogenase from *Ogataea polymorpha Op*GDH^31^ in the D452-2 strain of *S. cerevisiae* showed strong effects compared with native gene *Sc*GCY1 even if a *Sc*GCY1 integrated with other endogenous oxidative pathway genes (Glycerol proton symporter of the plasma membrane *Sc*STL1, *Sc*DAK1, *Sc*DAK2, and *Sc* TPI1) in recombinant strain GF2 (Table 1). The strain harboring the GDH gene, which named GDH, is consuming glycerol faster than GF2 with an increase of 21% in ethanol production, whereas it was only 10% in GF2 compared with the parental strain (Fig. 2). In full aerobic fermentation (1/10 liquid culture/flask volume) of mixed glucose and glycerol using GDH strain improved the glycerol consumption and ethanol production from 25% to 40% and from 21%–64%, respectively, before switching to the re-utilization of ethanol when compared with the previous semi-aerobic condition (Figs. 2 and 3). These results indicating the first step for the efficiency of glycerol fermentation should be through an effective GDH started here with an act of *Op*GDH. Furthermore, we confirmed that glycerol consumption was through the constructed DHA, where glycerol consumption has not significantly decreased after knocked-out the *Sc*GUT1 gene, which is the first gene in the G3P pathway (Fig. 2). Also, activating the genes of the G3P pathway (*Sc*STL1, *Sc*GUT1, *Sc*GUT2, and *Sc*TPI1) in a recombinant strain named GA2 (Table 1) did not impose significant improvement in the ethanol production (Fig. 2).

**Fig.2.**
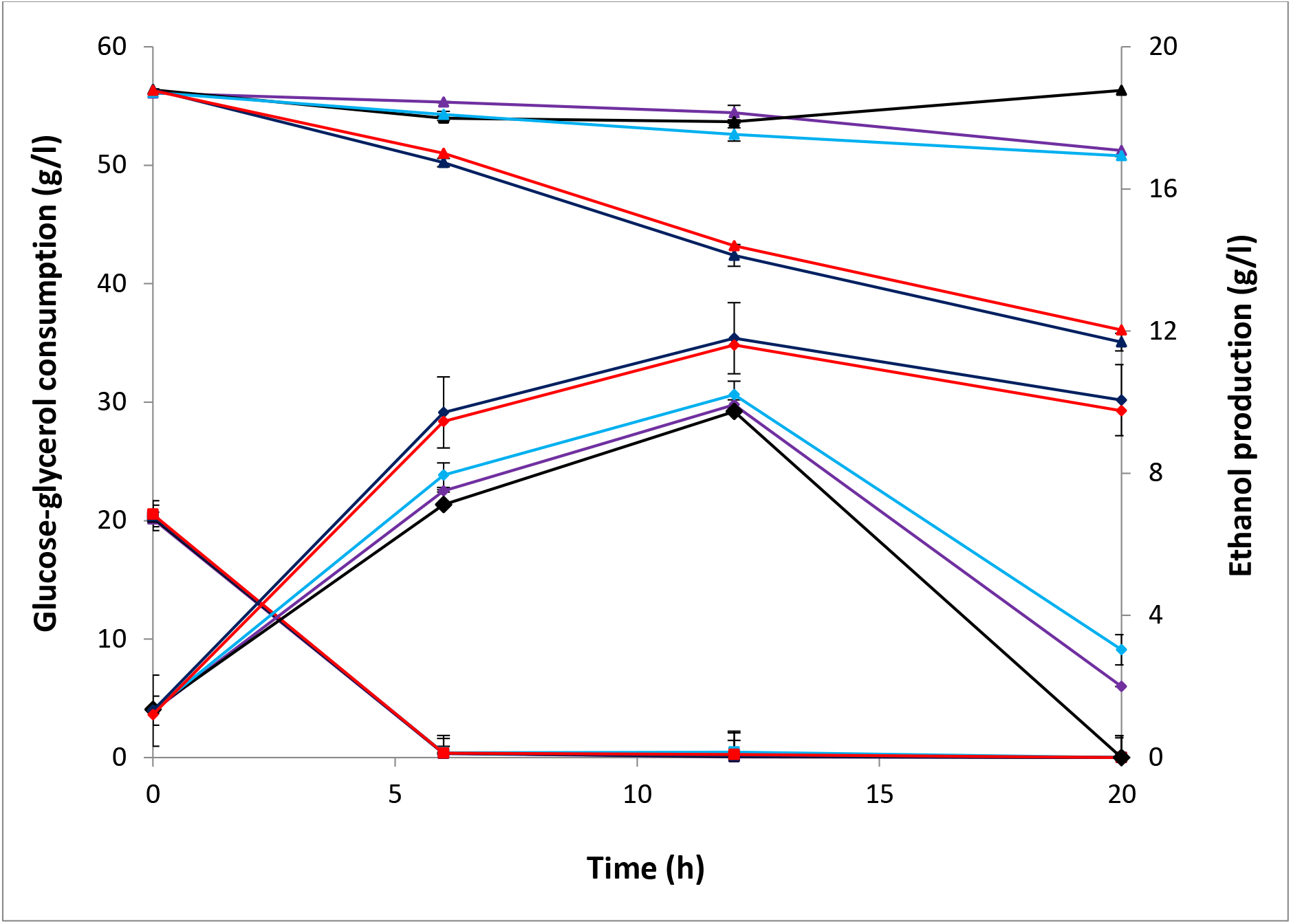
Time course for fermentation of glucose–glycerol by *S. cerevisiae:* ancestor (black lines colour); GF2 strain, overexpressing endogenous oxidative (DHA) pathways STL1, GCY1, DAK1, DAK2 and TPI1 (azure lines colour); GA2 strain, overexpressing endogenous assimilative (G3P) pathways STL1, GUT1, GUT2 and TPI1 (purple lines colour);GDH strain overexpressing glycerol dehydrogenase from *Ogataea polymorpha _Op_*GDH (blue lines colour); ΔGUT1-GDH (Red lines colour);glucose consumption (square symbols), glycerol consumption (triangular symbols) and ethanol production (rhomboid symbols). Data obtained from the mean of three independent experiments run at the time to decrease time-differences of sampling. Error bars represent the standard deviation of the mean and are not visible when smaller than the symbol size. We omitted the data indicated for fermenting the only YP due to the overlapped with the horizontal axis (YP in the medium was not converted to ethanol here).

**Fig.3.**
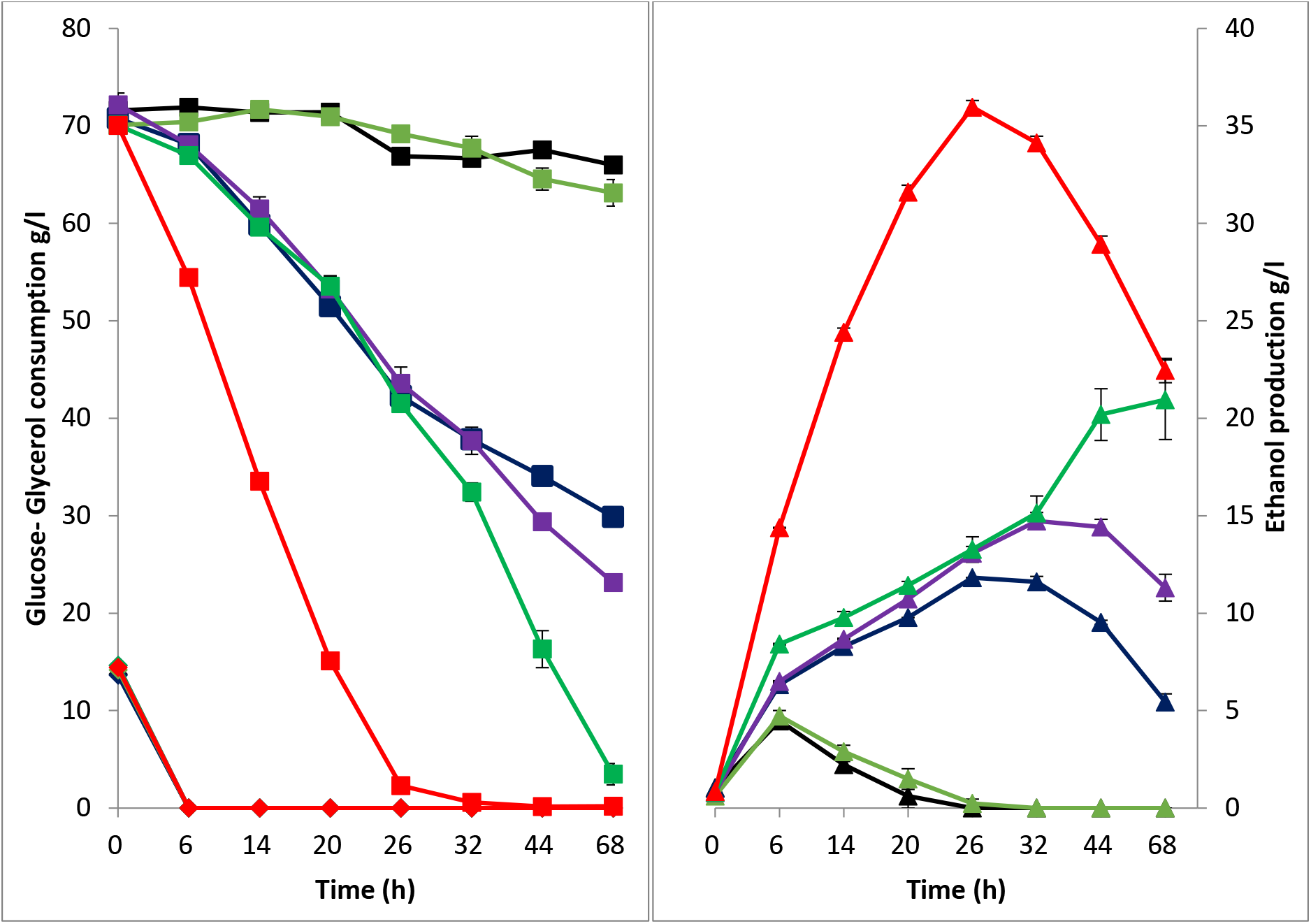
Comparison of the time course of glycerol–glucose fermentation between evolved *S. cerevisiae* strains during this study: ancestor, (black lines colour); NOXE/GPD1 strain (orange lines colour); GDH strain (blue lines colour); GDH-NOXE strain (purple lines colour); GDH-NOXE-FDT strain (green lines colour); (SK-FGG), GDH-NOXE-FDT-M1 (red lines colour); (A) Glucose consumption (rhomboid symbols), glycerol consumption (square symbols); (B) ethanol production (triangular symbols). Fermentation carried out in aerobic conditions in shaking flasks, 1/10 liquid culture/flask volume at 30°C with 180 rpm. Data obtained from the mean of three independent experiments run at the time to decrease time-differences of sampling. Error bars represent the standard deviation of the mean and are not visible when smaller than the symbol size. We omitted the data indicated for fermenting the only YP due to the overlapped with the horizontal axis (YP in the medium was not converted to ethanol here).

**Table 1:**
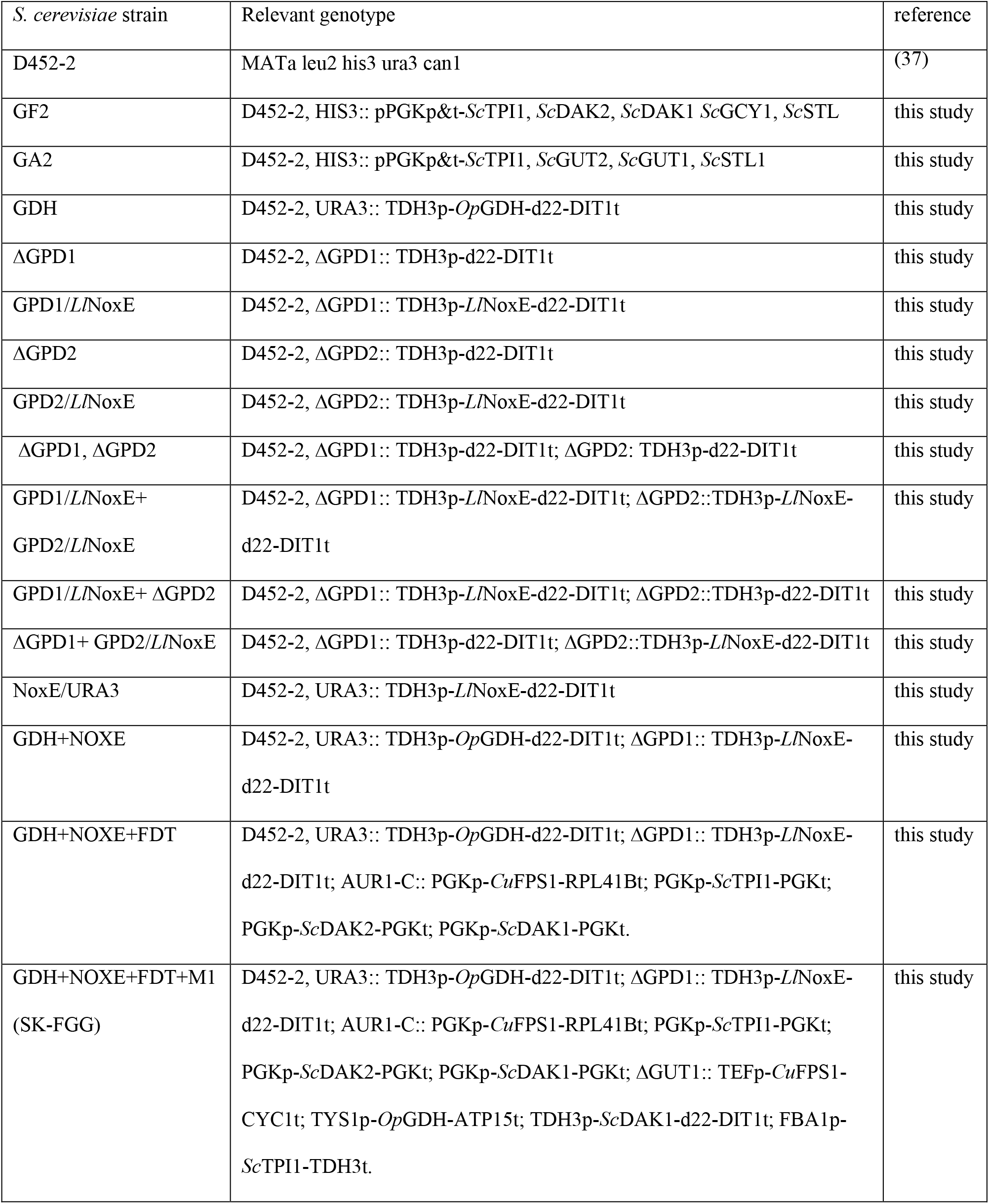
Characteristics of *S. cerevisiae* strains generated through this study

#### Step no. 2: efficient rewriting NADH pathway oxidation in *S. cerevisiae* by an O_2_-dependent dynamic shuttle of water forming NADH-oxidase (*Ll*NoxE) replaces GPD1

We comprehensively studied replacing GPD shuttles by water-forming-NADH-oxidase from *Lactococcus lactis* subsp. *lactis* (*Ll*NoxE). As the GPD shuttle is the first step in glycerol biosynthesis and represents one of the well-known systems for renovating a cytosolic NAD^+^ from NADH produces during the metabolic process such as those from the oxidation of glyceraldehyde 3-phosphate (GA3P). As a consequence, glycerol is secreting at the unbalances of redox-reactions toward NADH. Therefore, we first rated the participation levels of GPD1 and GPD2 in glycerol biosynthesis in our ancestor strain D452-2 at 10% of glucose fermentation by deleting each isoform separately. The data obtained, reveals that the participation ratio of GPD1 in glycerol biosynthesis was 82%, where glycerol secretion from Δ GPD1 was 0.47g/2.56g of wild type WT; while it was 23% (2.08g/2.56g WT) with Δ GPD2 (Table 2; Fig.S1a). Replacing GPD1 with *LlNoxE* reduced glycerol secretion by 98%, where 0.14g glycerol /2.56g of WT has secreted, whereas it only 29% with NoxE/ GPD2 strain which secreted 1.82g/ 2.56 of WT (Table 2; Fig.S1a). On the other hand, replacing both GPD1 and GPD2 with *Ll*NoxE not only prevent glycerol formation but also reduced glucose consumption significantly and obstructed cell growth and fermentation by almost the same levels at 15% while increased secretion of acetate by 2.46 fold (Table 2; Fig.S1a-e). Similarly, replacing GPD1 by *Ll*NoxE with deleting GPD2 (Table 2; Fig.S1a-e). Comparatively, the overexpressed *Ll*NoxE gene in the URA3 locus with the conserved native activity of glycerol biosynthesis pathway exhibited a moderate reduction in glycerol production of only 41% (1.53g/ 2.56g) in the D452-2 strain (Table 2; Fig.S1a). Notably, replacing GPD with *Ll*NoxE switched the glycerol production to an increase in acetate production (Table 2; Fig.S1b). Exclusively, replacing GPD1 with *Ll*NoxE is an excellent approach to eliminating glycerol formation during glucose fermentation to ethanol, where ethanol production increased by 9 % (0.474/ 0.432 g ethanol / g glucose as calculated in table 2, therefore, consolidating this replacement with GDH could improve glycerol conversion to ethanol.

**Table 2.** **shows;** glycerol secretion, acetate accumulation, ethanol production, ethanol yield rate, total glucose consumed and maximum growth (optical density OD) 20 h of ferment 10 % glucose as a sole provided carbon to YP medium under semi-aerobic conditions by the recombinant strains studied here for rewriting NADH cycles by water-forming NADH oxidase. Errors represent the deviation of the mean. Semi-aerobic (shaking flasks with 1/5 liquid culture/ flask volume at 150 rpm). * Not detected.

| Relevant strain | Glycerol secreted (g) | Acetic secreted (g) | Ethanol produced (g) | Ethanol yield g/ g glucose | Consumed glucose (g) | Optical density (OD) |
| --- | --- | --- | --- | --- | --- | --- |
| WT | 2.56±0.06 | 1.46±0.06 | 42.67±0.53 | 0.431±0.54 | 99.01±1.660 | 13.84±0.21 |
| ΔGPD1 | 0.47±0.07 | 1.66±0.07 | 45.78±1.09 | 0.455±1.11 | 100.58±2.20 | 18.10±0.21 |
| ΔGPD2 | 2.08±0.05 | 1.87±0.05 | 44.74±0.88 | 0.446±0.90 | 100.18±1.10 | 17.55±0.20 |
| ΔGPD1+ ΔGPD2 | 0.10±0.01 | 1.85±0.03 | 43.04±0.93 | 0.436±0.95 | 98.81±0.630 | 16.81±0.17 |
| NoxE/GPD1 | 0.14±0.01 | 2.81±0.08 | 47.92±0.93 | 0.474±0.96 | 100.97±1.33 | 16.22±0.19 |
| NoxE/GPD2 | 1.82±0.05 | 1.96±0.06 | 44.27±0.78 | 0.43±0.810 | 102.34±1.24 | 16.53±0.27 |
| NoxE/GPD1+ NoxE/GPD2 | ND* | 3.59±0.11 | 36.24±0.60 | 0.429±0.62 | 84.36±2.020 | 11.71±0.19 |
| NoxE/GPD1+ ΔGPD2 | 0.11±0.01 | 2.60±0.09 | 35.25±0.61 | 0.437±0.63 | 80.7±2.2530 | 13.04±0.18 |
| NoxE/GPD2+ ΔGPD1 | 0.11±0.02 | 2.77±0.06 | 43.25±0.86 | 0.435±0.87 | 99.28±2.110 | 13.73±0.19 |
| NoxE/URA3 | 1.53±0.06 | 2.93±0.06 | 43.40±0.77 | 0.43±0.820 | 99.7±1.3230 | 18.43±0.13 |

#### Step no. 3: integrating GDH and NoxE with ΔGPD1

Owing to the previous results from the recombinant GDH strain and the data of replaced the shuttles of oxidizing the cytosolic NADH by *Li*NoxE, therefore, we studied the recycling outputs of NAD^+^/NADH between the GDH and *Li*NoxE. In addition to this recycling, deleting GPD1 during that substitution with *Li*NoxE will abolish glycerol formation and decreases the ramification of DHAP, which consolidates the straightforward to glycolysis route. As a result of that thinking, we engineered a further strain that combined GDH with *Li*NoxE with a Δ GPD1 as listed in table 1. This round of recombination (GDH+NOXE strain) has tested for ability fermenting glycerol in comparison with GDH or *Li*NoxE, as well as the wild type strain. This innovative integration clearly showed improvements in both efficiency of glycerol conversion to ethanol and delayed in the time of reprogramming the cell to utilize a produced ethanol. In figure 3 at 6h, both strains of the ancestor, and the engineered GPD1/*Ll*NoxE started their re-utilize the produced ethanol from glucose without significant consumption in glycerol, where maximum ethanol produced was 4.7 g/l ethanol. In GDH strain, the time of reusing the produced ethanol delayed to 26h with raises in the ethanol production to 11.82 g/l, which represents 0.27 g ethanol/ the consumed glucose and glycerol. In the case of GDH+NOXE, the integration here not only boosted the ethanol production to 13.27 g/l (0.31g ethanol/ the consumed glucose and glycerol) at 26h, but also extended the fermentation time to 32h, and further raised production of ethanol to 14.42 g/l before the switching to consume that ethanol (Fig. 3).

#### Step no. 4: overexpressing the rest of the DHA pathway genes; TPI1, DAK1, DAK2, and FPS

Although clear impacts of recycled inputs in the previous recombination, we deduced a further limiting in the activity of other genes in the DHA pathway TPI1, DAK, FPS genes, which affect that full traffic of glycerol conversion to ethanol. Therefore, we proceeded to overexpress the rest of the genes included in the DHA pathway at this stage of systematic engineering. A promoter phosphoglycerate kinase (PGK) with its terminator had used to activate the endogenous genes TPI1, DAK1, and DAK2. The gene of glycerol facilitator from *Candida utilis* (*Cu*FPS1)^32^ had heterologous-expressed under the control of PGK promoter and Ribosomal 60S subunit protein L41B terminator (RPL41B). The previous recombinant-strain GDH+NOXE has used as a competent cell for receiving this one-set of genes in AUR-1C locus to generate a new strain, which had named GDH+NOXE+FDT (Table 1). Unequivocally, this fourth step of recombination solved one of the main problems in this study, where is prevented the phenomena of the switching to utilizing ethanol before the full consumption of glycerol. A consumption rate reached 1 g l^-1^h^-1^ and produced 20.95 g/l of ethanol by this recombinant strain. Nonetheless, its conversion efficiency of ethanol production appeared to be less than 48% of the theoretical value (Fig. 3).

#### Step no. 5: Super-expressing the DHA pathway genes by another copy of genes; *Sc*TPI1, *Sc*DAK1, *Op*GDH, and *Cu*FPS1 with abolishing the native G3P pathway

The stemming results from the fourth step of genetic engineering posited the effect of the limited activities of the other genes on paced productivity with the visibility to strengthen the pathway activity by another copy. We carefully selected and designed the strongest-expression systems that may not be affected by the repressors of regulators to constitutively-express this assortment of genes^33–36^. By merits of using the hybrid of Gibson assembly and PCR, we constructed one module named M1 (Table 3; Fig. S2) were their expression systems; TEF1 promoter-CYC1 terminator, TYS1 promoter-ATP15 terminator, TDH3 promoter-mutated d22DIT1 terminator, and FBA1 promoter-TDH3 terminator, respectively with genes *Cu*FBS1, *Op*GDH, *Sc*DAK1, and *Sc*TPI1. Therefore, we intensified a whole glycerol oxidation pathway by integrating another copy of genes *Cu*FPS1, *Op*GDH, *Sc*DAK1, and *Sc*TPI1 during the replacement of the GUT1, which abolished the G3P pathway, for continuing the overcoming of the previous inadequacies in this fifth stage of recombination. Interestingly, we griped unique findings with step five of recombination in the glycerol consumption and ethanol production that never reported in any organism with the evolved strain GDH-NOXE-FDT-M1 named SK-FGG (Table 1, Fig.3). Its consumption rate reached 2.6 g 1^-1^ h^-1^ from glycerol at the described experimental conditions, and the productivity paced 1.38 g l^-1^ h^-1^ of ethanol with conversion efficiency reached 0.44g ethanol/g glucose and glycerol (Fig. 4).

**Fig.4.**
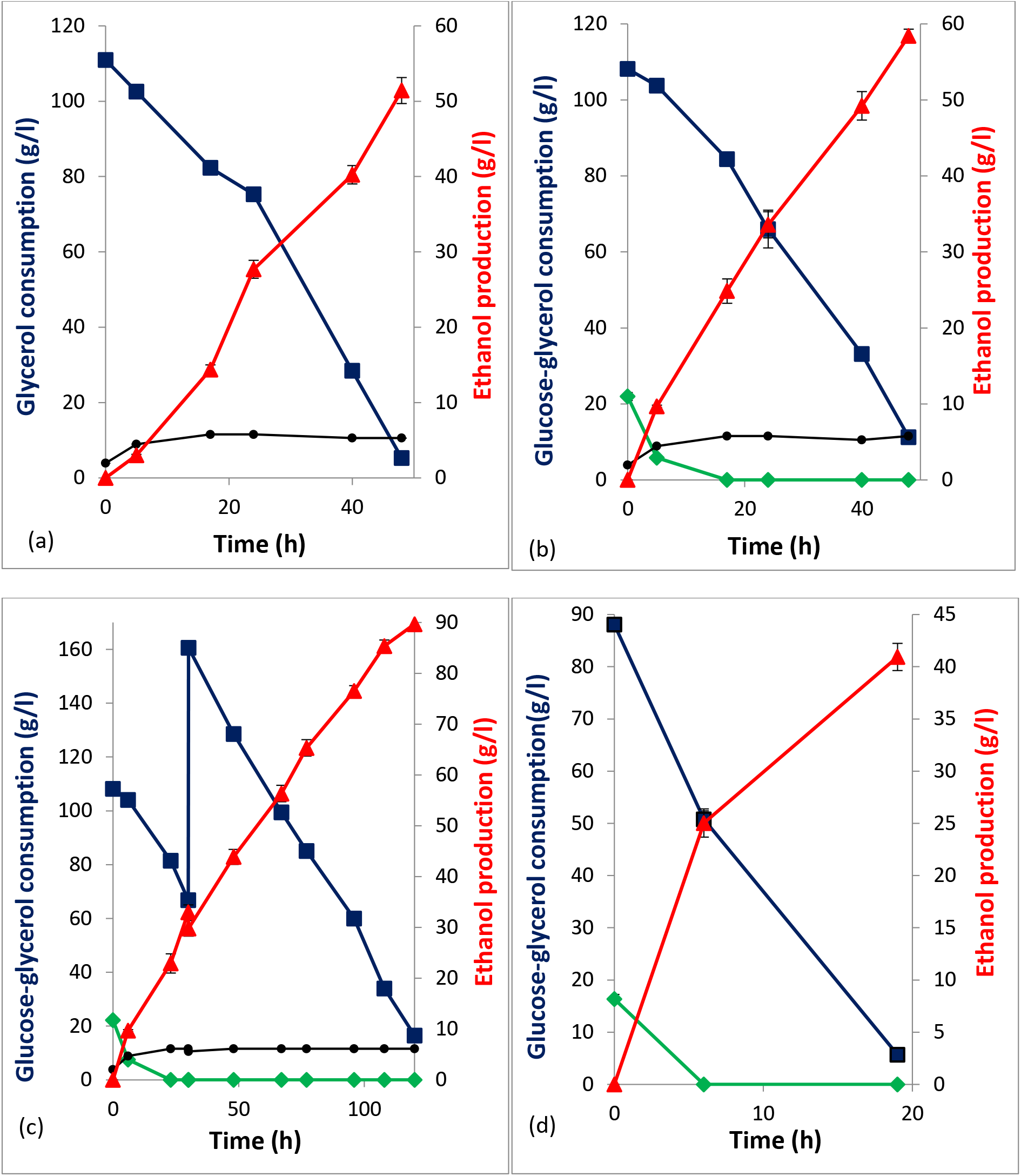
Time course of aerobic fermentation by glycerol fermentation of recombinant strain SK-FGG: glycerol consumption (blue lines colour with square symbols); Glucose consumption (green lines colour with rhomboid symbols); Ethanol production (red lines colour with triangular symbols); cell density (OD600) (black lines colour with circle symbols); (A) sole glycerol. (B) Mixed glucose and glycerol. (C) Mixed glucose and glycerol with glycerol fed-batching. Fermentation carried out in aerobic conditions in shaking flasks; 1/10 liquid culture/flask volume for (A), (B) and (C) and 1/30 liquid culture/flask volume for (D) at 30°C with 200 rpm. Data obtained from the mean of three independent experiments run at the time to decrease time-differences of sampling. Error bars represent the standard deviation of the mean and are not visible when smaller than the symbol size. We omitted the data indicated for fermenting the only YP due to the overlapped with the horizontal axis (YP in the medium was not converted to ethanol here).

**Table 3:**
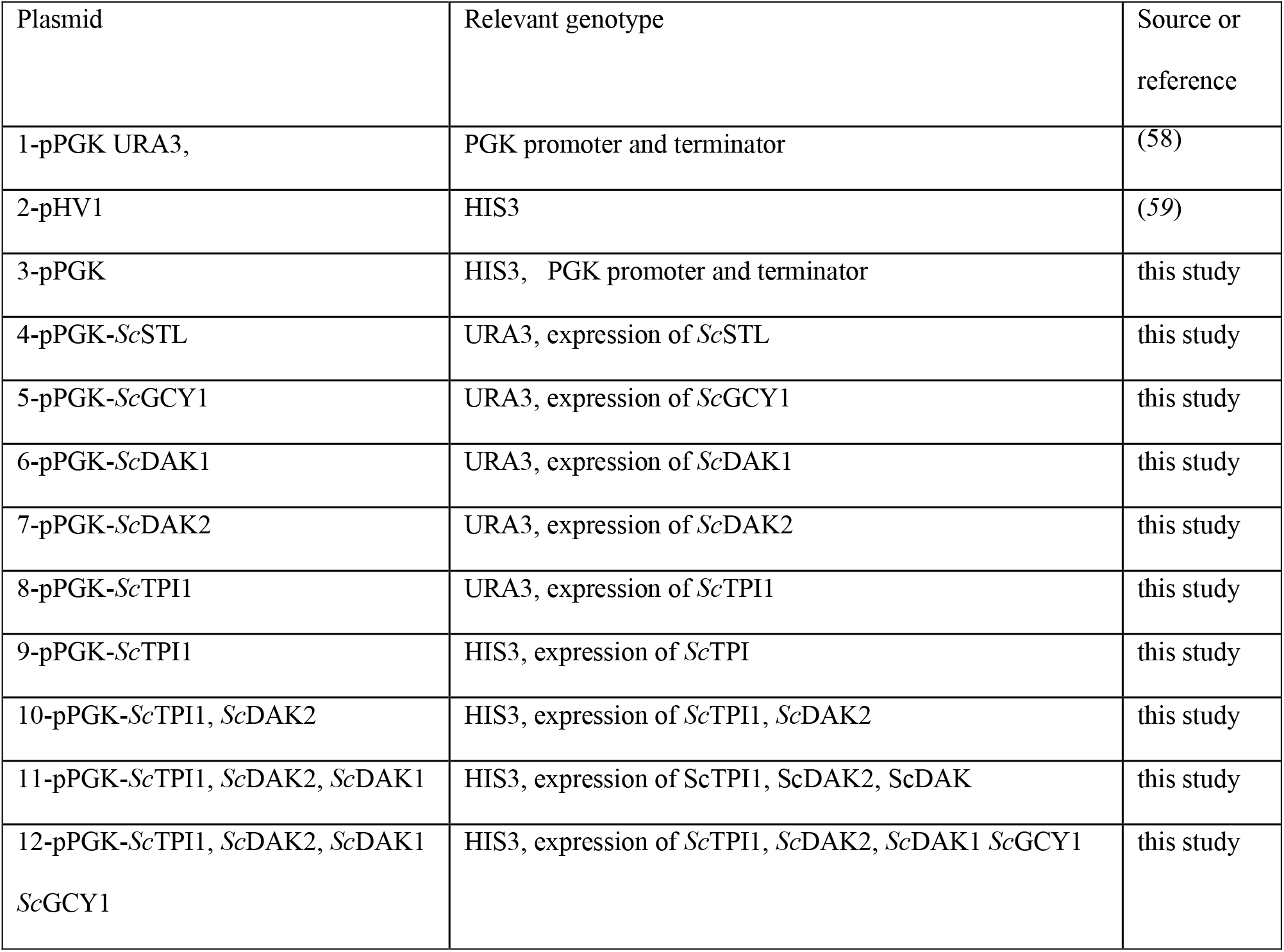
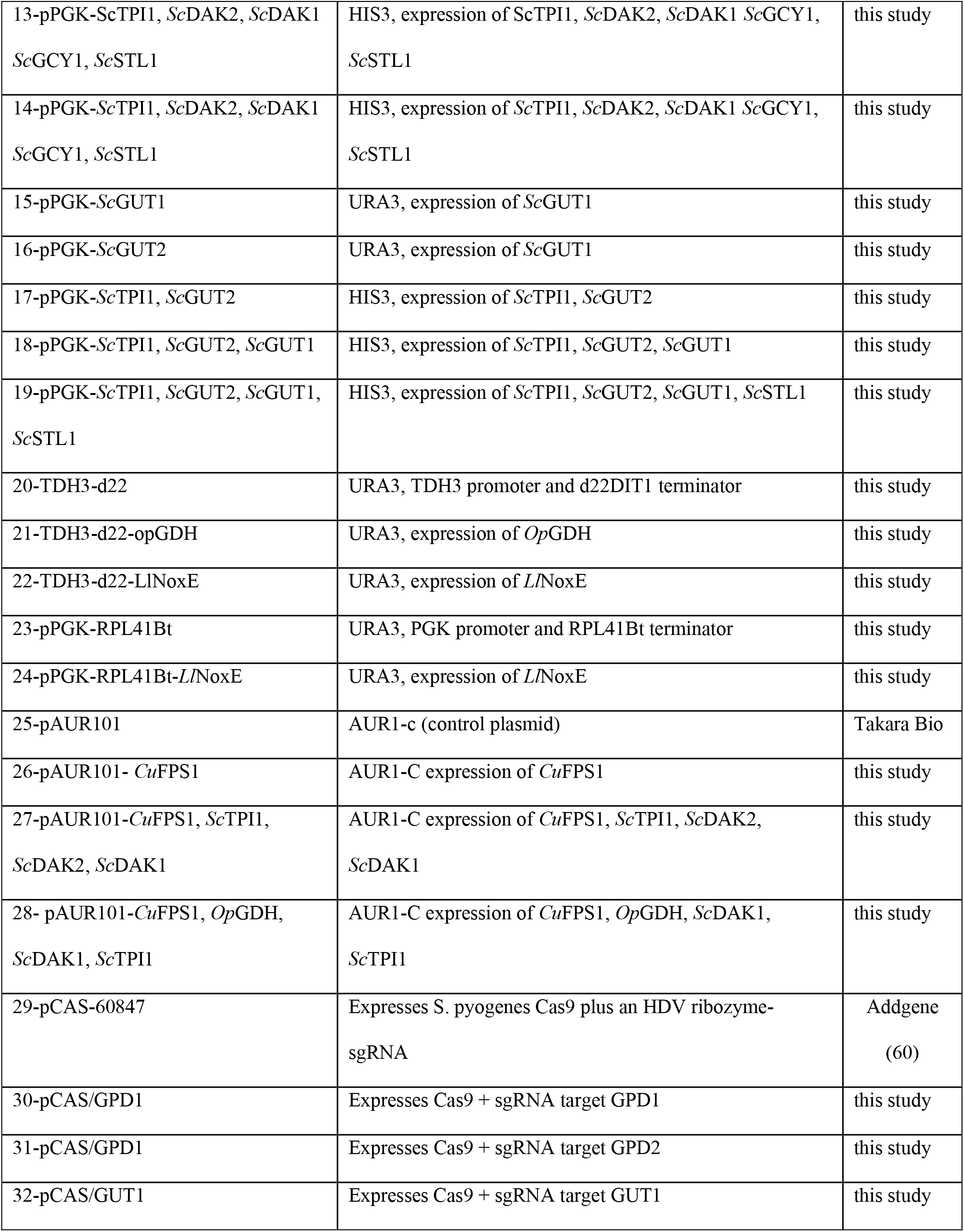
Constructed plasmids used in this study

### Osmotolerance of the engineered strain (SK-FGG) and the effect of higher aeration

the strain SK-FGG exhibiting outstanding performance in aerobic conditions at that higher initial concentration of glycerol in YP medium, where its conversion efficiency reached 0.49 g ethanol/g glycerol with a production rate of> 1 g l^-1^ h^-1^ of ethanol (Fig. 4a). Even with the mixing of glucose with glycerol at the same initial concentration, its conversion efficiency was comparatively the same (Fig 4b). Interestingly, the strain engineered here glows in its capacity to harmonize fermenting the glycerol with glucose, as well as, accumulation of 9% of bioethanol with additional fed-batching of glycerol, although the efficiency decreased to 0.43 g ethanol/g glycerol (Fig. 4c). Notably, increasing the aeration by increasing the volume of flasks with keeping the constant of the broth volume accelerated the glycerol consumption remarkably to >5 g l^-1^ h^-1^. Also, the rate of ethanol production increased to >2 g l^-1^ h^-1^. Nonetheless, its conversion efficiency decreased to 0.42 g ethanol/g glycerol (Fig. 4d). We observed some other minor uncharacterized peaks during the analysis of these samples, as a further point for research in the future. It is also worth mentioning that a strain SK-FGG has proved its capability to convert glycerol at larger volumes where we scaled up the experimental capacities to 1, and 3 liters via the mini-jar 5L fermentor (see Methods section). Nonetheless, its rates and efficiencies decreased due to higher fluctuating of the dissolved oxygen during the fermentation by our available system, where a more advanced control system is required.

## Discussion

Recently, microbial technologies for exploiting glycerol as a carbon source for producing valuable products have gained higher attention, where a considerable amount of glycerol as an unavoidable by-product from the expansion of biodiesel industries had accumulated. In our other scenario, glycerol has evidenced as a delignifing agent during pretreating biomass with alum in the glycerolysis process^30^. Therefore, working on engineering the yeast genetically for generating the ability to convert glycerol into ethanol becomes inevitable for such all of these perspectives. *S. cerevisiae* has full genes for two metabolic pathways (DHA and G3P) for glycerol catabolism showed in Fig.1; nonetheless, glycerol had considered as non-fermentable and unfavorable carbon as a feedstock^5, 15^. The distinctive differences seen in the ability to grow on the synthetic medium had based on the genetic background of strains. Besides, that growth is a quantitative trait based on alleles on genome^21^. Hence, prompted to scrutinize in the strain used here, where D452-2 originated from three different ancestor strains via five sequential segregates crossing, one of the parental strains is known its belonging to *S. cerevisiae* S288C^37–39^ (Table 1 and Fig. S3). D452-2 didn’t show the ability of growth on synthetic medium without supplementing supports of uracil, leucine, and histidine, although we confirmed that the UBR2 allele not truncated as in the CEN.PK family^22, 23^. At the moment, it is not clear whether this disability to grow on synthetic medium in our strain related to the genetic background as those of the negative-glycerol grower in 13 of *S. cerevisiae* strains^21^ or generated by the interrupted genes Ura3, Lue2, and His3 in D452-2 strain. Further studies are needed to reveal this point or ranking its growth rate with that previous well studied strains^21^ to quantify trait the growth alleles on synthetic medium. In addition to these unstudied points, we decided to use the rich medium represented in yeast peptone (YP), where it showed a significant accumulated amount of ethanol onset fermentation of glycerol with engineered for production of 1, 2-propanediol^26^.

Although revoking the TPI1 gene has considered a pivotal hub for the production of glycerol from glucose^40^, which is the reverse direction here, it hasn’t integrated with the previous study that examined overexpressing the native DHA-pathway^28^. Therefore, we combined the overexpression of the TPI1 gene with the DHA pathway to track the restrictions in that oxidative pathway in fermenting glycerol. Hence, we constructed a strain named GF2, which overexpressed its genes *Sc*STL1, *Sc*GCY1, *Sc*DAK1, *Sc*DAK2, and *Sc*TPI1. Concurrently, we recognized the limited activity of the *Sc*GCY gene compared with a glycerol dehydrogenase from *Ogataea polymorpha*^41^. Therefore, we constructed a yeast harbored *Op*GDH named GDH to be testing with GF2 during fermenting glycerol. As a result of these comparing studies, such an active *Op*GDH gene is the first key for deciphering glycerol fermentation, although the sole integration of *Op*GDH not enough to induce an efficient fermentation (Fig. 2). On the other hand, overexpressing the native glycerol catabolic pathway G3P in strain named GA2 did not demonstrate promising results as this oxidative pathway. The assumption that may be contemplating here is the limit of the respiratory chain during glycerol consumption, thus, restrict the renovation of FAD^+^ for converting glycerol 3-phosphate to DHAP through GUT2, considering this phosphorylated G3P-pathway is subjecting to the repression and transcriptional regulation with the presence of glucose^17–20, 42, 43^.

Besides the induction for expressing target genes, one of the other main obstacles affecting the efficiency of microbial production is to meet the stoichiometries of the engineered metabolic pathways, cofactors, and ATP/oxygen ratios, especially those pathways which require cofactors for their activation^2, 9, 44^. As long as an integrated *Op*GDH in *S. cerevisiae*, we visualized the GPD shuttles will promote to activates the oxidation of the plethora from cytosolic NADH and a reduction of DHAP into G3P pathway. Besides, a ramification of glycerol 3-phosphate into glycerolipid pathway takes place^45^. Likewise, the DHAP may be distributing into phospholipid and methylglyoxal biosynthesis^45, 46^. Moreover, inasmuch of fermentable sugar, especially in the presence of oxygen, there is a plethora of cytosolic NADH. As a result, there is a need for shuttles for re-oxidizing this surplus. The shuttle of GPD plays an essential role in this regard with reducing DHAP to glycerol 3-phosphate to keep this homeostasis. Intracellular redox homeostasis in *S. cerevisiae* comprising > 200 reactions; thus, the shuttles oxidizing NADH has been well studied^16, 44, 47^ One of the interesting ones is the catalyzing oxidation of cytosolic NADH by heterologous-overexpressing a water-forming oxidase gene from *Streptococcus pneumoniae* in *S. cerevisiae* for reducing the cytosolic NADH, and the overflow to glycerol biosynthesis^44^. Nonetheless, there are no studies regarding the effects of replacing the native shuttles of GPD by other shuttles for oxidizing the cytosolic NADH, such as those of water-forming. Therefore in the second round of recombination in this study, we comprehensively focused on preventing that overflows to glycerol biosynthesis with the conservation of intracellular redox homeostasis during fermenting glucose. The Investigation of the nine constructed strains of either deleted or replaced GPD1 or/and GPD2 by the NoxE gene, showed replacing GPD1 by *Li*NoxE is the best approach, where glycerol biosynthesis effectively abolished by 98%, and an improvement in the fermentation efficiency by 9% (Table 2).

Expectedly, this single replacement will not exhibit further progress toward glycerol fermentation (Fig.3). Assuredly, we referred to the act of the low activity of native glycerol dehydrogenase ScGCY1^41^. As detailed above, a ramification of DHAP represents another hindrance for the straightforward toward glycolysis from glycerol. In this juncture, a reduced circulation of DHAP into the G3P pathway had confirmed to be efficient for glycerol fermentation by integrating this replacement of GPD1 by *Li*NoxE within the GDH strain. As expected, the strain harbored this unique point of integration (GDH-NOXE) in this regard showed substantial improvement in ethanol production from glycerol reached 28 % compared with GDH strain at that studied conditions, which not considered the other parameters such as oxygen level (Fig.3). The role of abolishing GPD1 had explicitly calculated from the data of fig. 3, which has represented 43 % of that improved ratio. Utilize the recycles of cofactors NADH/NAD^+^ for production of 1, 2-propanediol has been well studied during fermenting glycerol^26^. Nevertheless, it seems non-stoichiometries of cofactors in the engineered pathway have compensated with the flowed to the ethanol accumulation relatively with rich media and the faster growth rates at the onset of fermentation and lately with re-consumption of ethanol by alcohol dehydrogenase (ADH2)^26^.

The importance around the activation of the other genes in the DHA pathway has confirmed, through the continued bioethanol production until the full consumption of glycerol (Fig.3). Although we didn’t evaluate the effect of overexpressing each gene individually, we recognized the cooperative effects for overcoming that traditional-ambiguous phenomenon of re-consuming the onset produced ethanol earlier than the full consumption of glycerol. In this regard, it had reported that the permeability of the three-carbon compounds including glycerol in *Candida utilis* is much faster than in the baker’s yeast, which supports the efficient utilization of glycerol, even at low concentrations^48^. Therefore, heterologous-expressing *Cu*FPS1 in *S. cerevisiae* could support the influxes of glycerol in our strain as reported earlier^25, 26, 32^ Also, DAK1 and DAK2 had characterized for detoxifying DAH, with *K*m_(DHA)_ of 22 and 5 μM and *K*m(ATP) of 0.5 and 0.1 mM, respectively, thus overexpressing DAK2 which is a much lower *K*m_(DHA-ATP)_ with DAK1 here definitely detoxify DHA that may accumulate by the action of the introduced *Op*GDH and *Cu*FPS1 in this. Besides, efficiently transfer DHA to DHAP. Furthermore, with presented genetic modifications during the introduction of *Cu*FPS1, and *Op*GDH, with overexpressing *Sc*DAK1, and *Sc*DAK2, DHAP may be accumulated to substantial concentration to influx the G3P-pathway through the GPD2 or saturated the native activity of TPI1 to be turned into pentose phosphate pathway especially with the presence of glucose^49^. Through scrutinizes in the previous studies abolished the activity of TPI1, we recognized the pivotal role of overexpressing TPI1 in this study, where the intracellular concentration of DHAP accumulated to 30-fold^50^ and when this deactivation further coupled with other deletions of NDE1, NDE2, and GUT2, the fermentation product had shifted from ethanol to glycerol^40^.

However, integrate one copy of the whole DHA-pathway with NoxE generated the ability of yeast to convert all supplemented glucose and glycerol to ethanol. Nonetheless, we recognized that conversion efficiency may still be affected by the robustness of native programed-glycolysis. Thence, further strengthening of the whole genes in the DHA pathway by another copy under different expression systems could overcome this obstacle. Interestingly, the other copy of *Cu*FPS1, *Op*GDH, *Sc*DAK1, and *Sc*TPI1 that replaced GUT1 met our expectations and reaches by efficiencies and the production rates to that comparable with the industrial application, where the efficient conversions reached 98% of theoretical ratio with production rates 1.38 g l^-1^ h^-1^. A potential using the strategy of multi-copy with optimizing the stoichiometries of the metabolic pathway had considerably boosted the production, e. g. six copies of the farnesene synthase gene, which integrated into yeast to improve the synthesize of farnesene^2^. Here, with the second copy of integration, we further selected highly constitutive expressing system in yeast^33–36^ to extend the production levels and efficiencies, where TEF1 promoter-CYC1 terminator, TYS1 promoter-ATP15 terminator, TDH3 promoter-mutated d22DIT1 terminator, and FBA1 promoter-TDH3 terminator, respectively with genes *Cu*FBS1, *Op*GDH, *Sc*DAK1, and *Sc*TPI1. Owing to the efficient SK-FGG strain generated, and its introduced pathway, oxygen supplements were the limit. Surprisingly, fermentation rates doubled with increasing aeration to >2 g l^-l^ h^-l^. Nonetheless, we are currently working on further improvements to increase the efficiency during such production rates, as well as utilize glycerol’s high reduction merit for improving the fermentation efficiencies of other carbons.

In this study, we are reporting the discovery for the modeling of glycerol traffic to the industrial levels of bioethanol production. This modeling includes the integration of (i) Impose vigorous expression to all genes in the glycerol oxidation pathway DHA. (ii) Prevalence of the glycerol oxidation by an oxygen-dependent dynamic by water-forming of NADH oxidase NoxE, which controls the reaction stoichiometries with regenerate the cofactor NAD^+^. (iii) Revoking the first step of both glycerol biosynthesis and glycerol catabolism through G3P, as shown in (Fig. 1). Our study provides an advancing use of metabolic engineering for re-routing the glycerol traffic in *S. cerevisiae* with tracking ethanol production to the highest levels that never attained by any other native or genetically engineered organism^27, 28, *52 53 54*^. Enormous considerations for the global demands for bioethanol reported, although the limited resources. Thus, it constrained the global annual bioethanol production to nearly 28.5 million gallons, which represents <2.7% of the transportation fuels*^55–57^*. Therefore, the current study is expanding the horizon of utilizing the surplus of glycerol directly to produce bioethanol. By fermenting glycerol, we avoid the burden of the pretreatments, and enzymatic saccharification, besides the problems of fermentation inhibitors. Furthermore, the engineered strain in this study has revised a promising scenario of biorefinery. SK-FGG dramatically improved bioethanol production from bagasse with the incorporation of glycerol, which has pretreated the bagasse with alum^30^ and produced the industrial levels of bioethanol from that glycerolysis mixture (Data not showed). The outcome of this study is to promote the association between bioethanol and biodiesel industries, which may develop their expansions with overburdening the sustainabilities. It may also prevent a decrease in the present glycerol price as well as broadening the horizons of glycerol producing industries for the production of glycerol.

## MATERIALS AND METHODS

### Section I: Cassettes and plasmid construction in this study

#### 1- Construction of pPGK-*Sc*TPI1, *Sc*DAK2, *Sc*DAK1 *Sc*GCY1, *Sc*STL and pPGK-*Sc*TPI1, *Sc*GUT2, *Sc*GUT1, *Sc*STL1 plasmids

We obtained the genes’ DNA from the ancestor strain D452-2 to clone the plasmids in this section. At first, disrupting the cell walls by re-suspended toothpick-touched cells in 20 μl 30 mM NaOH at 95°C for 10 min and then used directly as a template for PCR, fresh 1μl of that disrupted cells is suitable for 50 μl of PCR mixture. All primers used to obtain the native genes were designed based on the sequences available on the Saccharomyces Genome Database (SGD): https://www.yeastgenome.org/. For assembling the following plasmids: pPGK-*Sc*TPI1, pPGK-*Sc*DAK2, pPGK-*Sc*DAK, pPGK-*Sc*GCY1, and pPGK-*Sc*STL, the following genes: STL1, GCY1, DAK1&2, and TPI1 obtained from genomic DNA of ancestor strain by PCR. High fidelity polymerization of KOD-plus neo with their corresponded primers (Section 1 – table S1) used during this amplification. Xhol site of DAK2 deleted before cloning. These DNA genes were purified from the PCR mixtures by columns obtained from Nippon Genetics Co., Ltd., with its accessories, and then form their cohesive ends according to the designated primers and restriction enzymes. At first, we separately cloned each gene in pPGK/URA3 plasmid^58^, under the control of the expression system PGK promoter and its terminator (Table 3). We further replaced the URA3 gene in a pPGK-URA3 plasmid with a gene of HIS3 ^(Ref59)^ using the feature of a synthetically adding an overlapped sequences from pPGK plasmid to HIS3 marker using PCR and primes and vice-versa (Section 3 – table S1). Then, a Gibson Assembly Master Mix assembles the overlapping ends of the two fragments to form PGK-HIS3 plasmid. With construct pPGK-HIS3 plasmid (Table 3), we granted HIS3 locus for homologous recombination in *S. cerevisiae* after linearizing the plasmid at the BsiWI site. We obtained the previous plasmids and confirming their genes sequences by sequencing, detailed relevant primers listed in (Section 2 – table S1). Next, we cut XhoI/SalI-TPI1 cassette and inserted it into XhoI/SalI sites of a newly constructed plasmid pPGK-HIS3 plasmid. Following, integrating the genes with their systems together in one plasmid started by connecting the DAK2 set into the SalI site of the template plasmid started here by pPGK-TPI 1. The deadly ligations (XhoI/SalI sites) that cannot reopen were used repeatedly during the ligation of new cassettes and form the new plasmids. Repeatedly, DAK1, GCY1, and STL1 combined. Ultimately, pPGK-*Sc*TPI1-*Sc*DAK2-*Sc*DAK1-*Sc*GCY1-*Sc*STL1 plasmid was constructed (Table 3). Continuing with the same procedures, the plasmid pPGK-*Sc*TPI1-*Sc*GUT2-*Sc*GUT1-*Sc*STL1 also established.

#### 2- Construct TDH3p-d22DIT1t, TDH3-d22-opGDH and TDH3-d22-LlNoxE plasmids

##### a. **Cassette1:** partial end of GPD1promoter-TDH3p-d22DIT1terminator-partial front side of GPD1 terminator and TDH3p-d22DIT1t plasmid

The mutated terminator d22DIT1t purchased from (Integrated DNA Technology (IDT) Company, Tokyo, Japan) according to the published sequences^35^. TDH3 promoter magnified from the genomic DNA of the ancestor strain D452-2 using PCR and the designated primers (Section 4 – table S1). All primers bought from FASMAC Company, Japan. Moreover, flanking sequences added upstream of the promoter and downstream of the terminator using the feature of PCR polymerization with primers possess a long-desired tail, and a further extension to those flanking sequences with the addition of restriction sites accomplished by PCR in the second step (Section 4 – table S1). Then, cohesive the ends of that couple of DNA fragments by restriction enzymes XhoI, NotI for the first fragment and NotI, SalI for the second one. After purification of the fragments using agarose gel and columns of Nippon Genetics Co., Ltd., one-step cloning coupled the TDH3 promoter and mutated DITI terminator into XhoI/SalI of PGK/URA3 plasmid. Then TDH3p-d22DIT1t-URA3 plasmid constructed (Table 3).

##### b. **Cassette 2:** partial end of GPD1promoter-TDH3p-*Op*GDH-d22DIT1t-partial front side of GPD1terminator and TDH3-d22-opGDH plasmid

The previously constructed TDH3p-d22DIT1t/URA3 plasmid used as a template for constructs the next plasmid by further cloning an *Ogataea polymorpha* glycerol dehydrogenase gene (*Op*GDH), deposited in gene bank under the accession number XP_018210953.1. *Op*GDH synthetically purchased from the IDT Company. Primers listed in (Section 4 of Table S1), and full sequences are available in (Table S2).

##### c. **Cassette 3:** partial end of GPD1promoter-TDH3p-*Ll*NoxE-d22DIT1t-partial front side of GPD1 terminator and TDH3-d22-*Ll*NoxE plasmid

We also purchased, (synthetically from IDT Company), the water-forming NADH oxidase gene of *Lactococcus lactis* based on sequence available on gene bank accession number AAK04489.1 and cloned it into TDH3p-d22DIT1t to assemble TDH3-d22-*Ll*NoxE plasmid (Tables 3, and S2).

##### d. **Cassette 4:** partial end of GPD2promoter-TDH3p-*Ll*NoxE-d22DIT1t-partial front side of GPD2 terminator and TDH3-d22-*Ll*NoxE plasmid

In this step, we replaced the flanking sequences of GPD1 promoter and terminator with GPD2 using PCR and the primers listed in (Section 5 – table S1)

#### 3- Construct multiplex pCAS-gRNA-CRISPR systems

The multiplex pCAS-gRNA system was a gift from Prof. Jamie Cate^60^ (Addgene plasmid # 60847; https://www.addgene.org/60847/). For that, we used the online tool for the rational design of CRISPR/Cas target to allocate the highest probability of the on-target sites for the gRNA in the genomic DNA of *S. cerevisiae:* https://crispr.dbcls.jp/ ^(Ref61)^. Accordingly, the sequence of the primers designed based on previously allocated sequence (20 bp before the PAM), with another 20 bp from sgRNA or HDV ribozyme for overlapping (Sections 4.2, 5.1 and 7.1 – table S1). First, PCR synthesizes two fragments from the template, pCAS-gRNA plasmid. The first one amplified by PCR using a forwarding primer called pCas For., which located upstream of the gRNA scaffold at the SmaI site of pCas and the antisense primer, which has a reverse sequence of target gRNA. The second fragment amplified by forwarding primer, which has a sense sequence of gRNA and a reverse primer called pCas Rev., located downstream of the gRNA scaffold (Section 4.2 – table S1). After purifying each DNA part, overlapping and integration carried out by PCR using the pCas For., and pCas Rev. primers. Then, the produced fragment restricted to SmaI-PstI sites for the cloning into a truncated pCAS-gRNA plasmid with SmaI-PstI. As a result, a new multiplex pCAS-gRNA plasmid formed. Repeatedly steps have done with constructing all multiplex pCAS-gRNA plasmids that target GPD1, GPD2, and GUT1 (Table 3). We confirmed the newly constructed systems by sequencing their whole scaffolds.

#### 4- Construct pAUR101-*Cu*FPS1 and pAUR101-*Cu*FPS1, ScTPI1, *Sc*DAK2, *Sc*DAK1 plasmids

*Candida utilis*, NBRC 0988, obtained from the National Biological Resource Center (NBRC) of National Institute of Technology and Evaluation NITE, Japan. It used as a template for getting the gene glycerol facilitator FPS1 (*Cu*FPS1). The sequence of *Cu*FPS1 included in the deposited gene bank accession number BAEL01000108.1. Original pAUR101 plasmid purchased from Takara Bio, Inc., Japan, and the primers used to establish this plasmid listed in (Section 6 – table S1). A full sequence for cassette, PGK-*Cu*FPS1-RPL41Bt, transferred to (table S2). First, we constructed a pAUR101-PGKp-RPL41Bt vector by one-step cloning of the SmaI-Not1 PGK promoter (fragment1) and NotI-SalI-RPL41B terminator (fragment 2) into SmaI-SalI pAUR101 vector and then cloning a cohesive ended-NotI-*Cu*FPS gene into dephosphorylated NotI site of pAUR101-PGK-RPL41B vector to assemble pAUR101-PGKp-*Cu*FPS1-RPL41Bt vector. To constitute pAUR101-*Cu*FPS1, *Sc*TPI1, *Sc*DAK2, *Sc*DAK1 plasmid, we detached the set of cassettes, *Sc*TPI1, *Sc*DAK2, and *Sc*DAK1, from previously constructed plasmids, pPGK-*Sc*TPI1, *Sc*DAK2 and *Sc*DAK1 (Table S1), using restriction enzymes Xhol-SalI and re-inserted that set of cassettes, *Sc*TPI1, *Sc*DAK2, and *Sc*DAK1, into the SalI site of pAUR101-PGK-*Cu*FPS1-RPL41B plasmid (Table 3).

#### 5- Construct Module M1; *Cu*FPS1, *Og*GDH, *Sc*DAK1, *Sc*TPI1 cassettes with flanking sequences of GUT1 promoter and terminator in plasmid pAUR101

At first, we obtained all fragments which will form the module M1 separately by PCR (Fig. S2); also, *Cu*FPS1and *Op*GDH genes mutated d22DIT terminator amplified from their synthetic DNA stocks, whereas other fragments magnified from the genomic DNA of the D452-2 strain (Fig. S2). The full sequence of the module M1 is also accessible in (table S2), and the details of the primer listed in (Section 7 – table S1). Purification of the 12 amplified DNA fragments was carried out on 1%–2% agarose gel and then recovered by the FastGene Gel/PCR Extraction Kit (Nippon Genetics Co. Ltd) according to the manufacturer’s protocol. Accordingly, we obtained highly purified fragments before the onset of assembles using the Gibson Assembly Master Mix. Effectively, we joined the first three parts seamlessly, as well as for every next three fragments (Gibson’s protocol). Also, we directly amplified each set by PCR and then purified them again on the agarose gel. Repeatedly, we gathered the first six parts, as well as the other six fragments, and then assembled the whole module M1. We further added the SacI site to the upstream of the module M1and SmaI site to the downstream as well. These restriction sites provided for cloning the module M1 into SacI-SmaI sites of pAUR101 vector to form pAUR 101-M1 (table 3). Finally, we transferred that vector, pAUR 101-M1, into *E. coli* as described previously and also confirmed the correct structure of M1 by sequencing the whole module M1 from pAUR-M1.

### Section II: Transformation and strains recombination in this study

All the previous plasmids stored in *E. coli* NEB 10-beta, for further the uses of production the required plasmids or cassettes, using the heat shock method according to the procedures provided with the competent cells. All plasmid extractions performed using the QIAprep Spin Miniprep Kit following the manufacturer’s protocol. All measurements of DNA were estimated using BioSpec-nano (Shimadzu, Japan) and stocked in freezing at −20 °C for future uses. Yeast transformation by Fast Yeast TransformationTM kit (Takara Bio) used for integrated linear pAUR101 vector and its associated genes in AUR1-C locus, linear pPGK plasmid with its cloned genes in either HIS3 or URA3 loci as well^37^. For achieving editing the genome and the replacement of GPD1, GPD2, and GUT1 genes with its designated DNA repairing cassette or module, we used the protocol of CRISPR-Cas9 genome engineering in *S. cerevisiae* cells^62^. We confirmed target replacements using PCR check for the inserted repairing cassettes with primers from upstream and downstream of the flanking recombined loci. Primers listed down in each section (Table S1). Furthermore, we cultivated up to 10 generations of the selected evolved strains to confirm the loss of pCAS plasmid and to re-confirm the recombination. All recombination strains and their genotypes were listed (Table 1).

### Section III: Fermentation procedures and analysis

The initial fermentation experiments tested in 100 ml shaking flasks with 1/5 (liquid/flask volume) with 200 rpm at 30°C for the estimation of semi-aerobic conditions while 1/10 (liquid/flask volume) for the aerobic conditions and then enlarged to 500 ml flasks. Additionally, we tested the scale-up of fermentation volume to 1L, 3L, using a mini jar 5L fermentor (TSC-M5L; Takasugi Seisakusho, Tokyo, Japan) equipped with a DO controller (DJ-1033; ABLE Corporation, Tokyo, Japan). The dissolved oxygen was adjusted automatically by the rotation speeds. Cells initially harvested for fermentation with the same volume of pre-culture YPD medium for ~24 h. The harvesting carried out by centrifugation at 6000 × g for 5 min at 4°C and washed with sterile water, then collected cells were re-supplemented by the Yeast-Peptone (YP) medium with glucose, glycerol or both, as shown in Figs. 2-4. Different initial concentrations tested to determine fermentation abilities at those different initial concentrations as well as with the fed patch for an estimate the maximum tolerance to the product under these unprecedented fermenting conditions. The cell density monitored using spectrophotometry at 600 nm (AS ONE, China).

All analyses estimated using auto-sampling a 10 μl to fractionated in an Aminex HPX-87H column (Bio-Rad Laboratories, Hercules, CA, USA), analyzed in a refractive index detector (RID-10A; Shimadzu) equipped to auto-sampled Ultra-Fast Liquid Chromatography (UFLC) (Shimadzu, Japan). Fractionation accomplished with 0.6 ml/min of a mobile phase 5 mM H_2_SO_4_ at 50°C. Reactant concentrations were estimated by monitoring the peak areas compared with the standards of the authentic reactant’s glucose, glycerol, ethanol, and acetic acid.

## Supplementary Materials

**Fig. S1.**
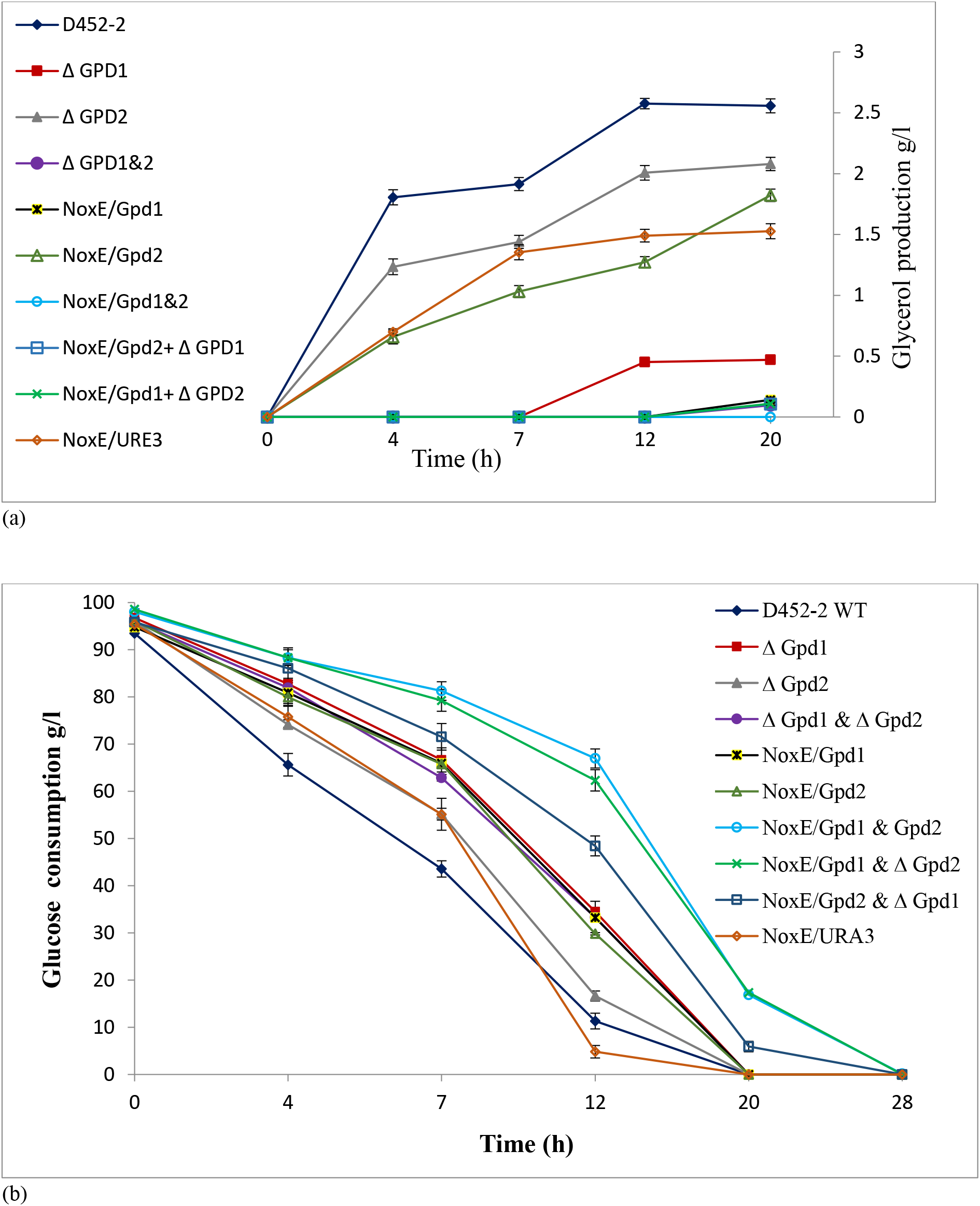
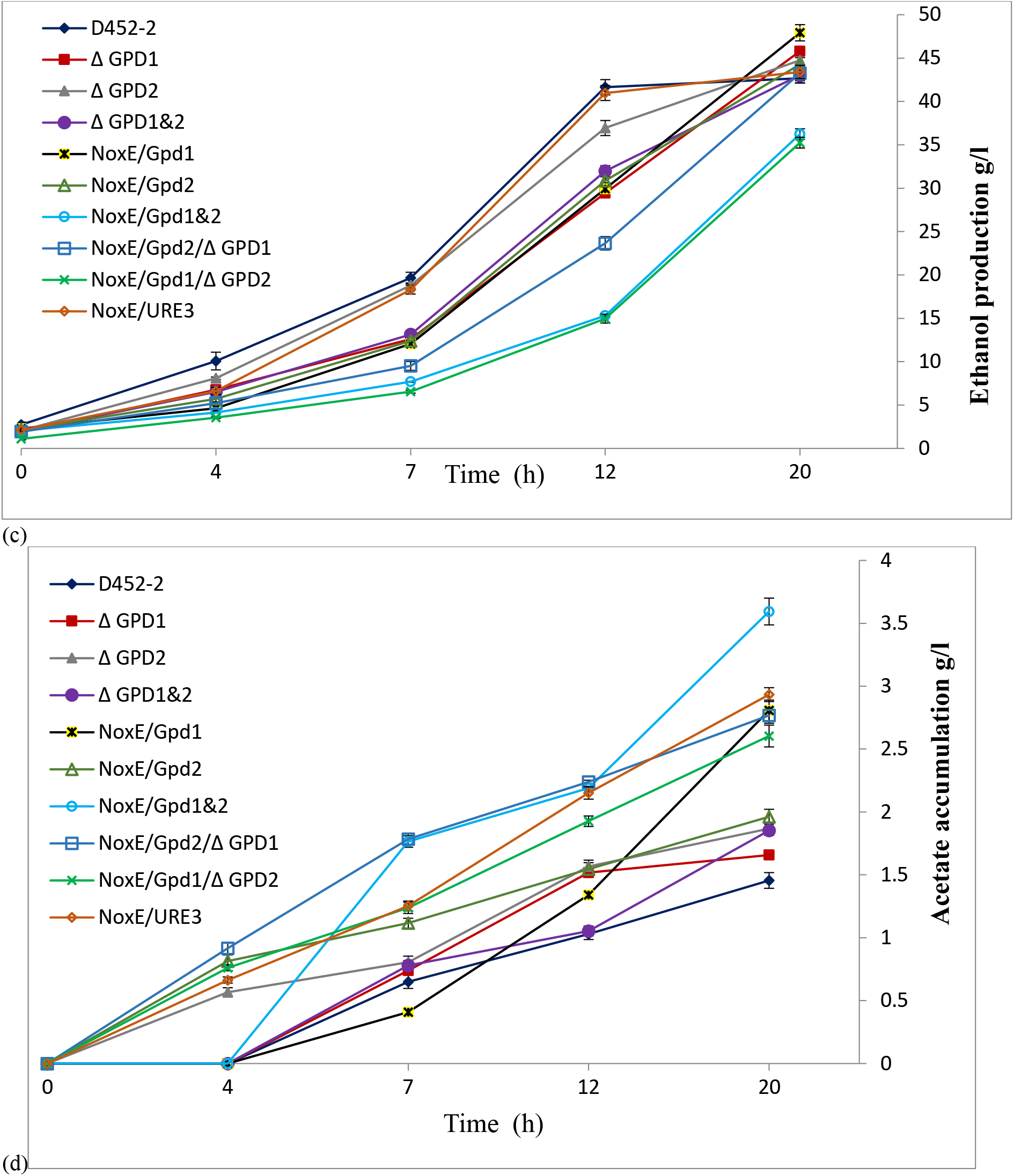
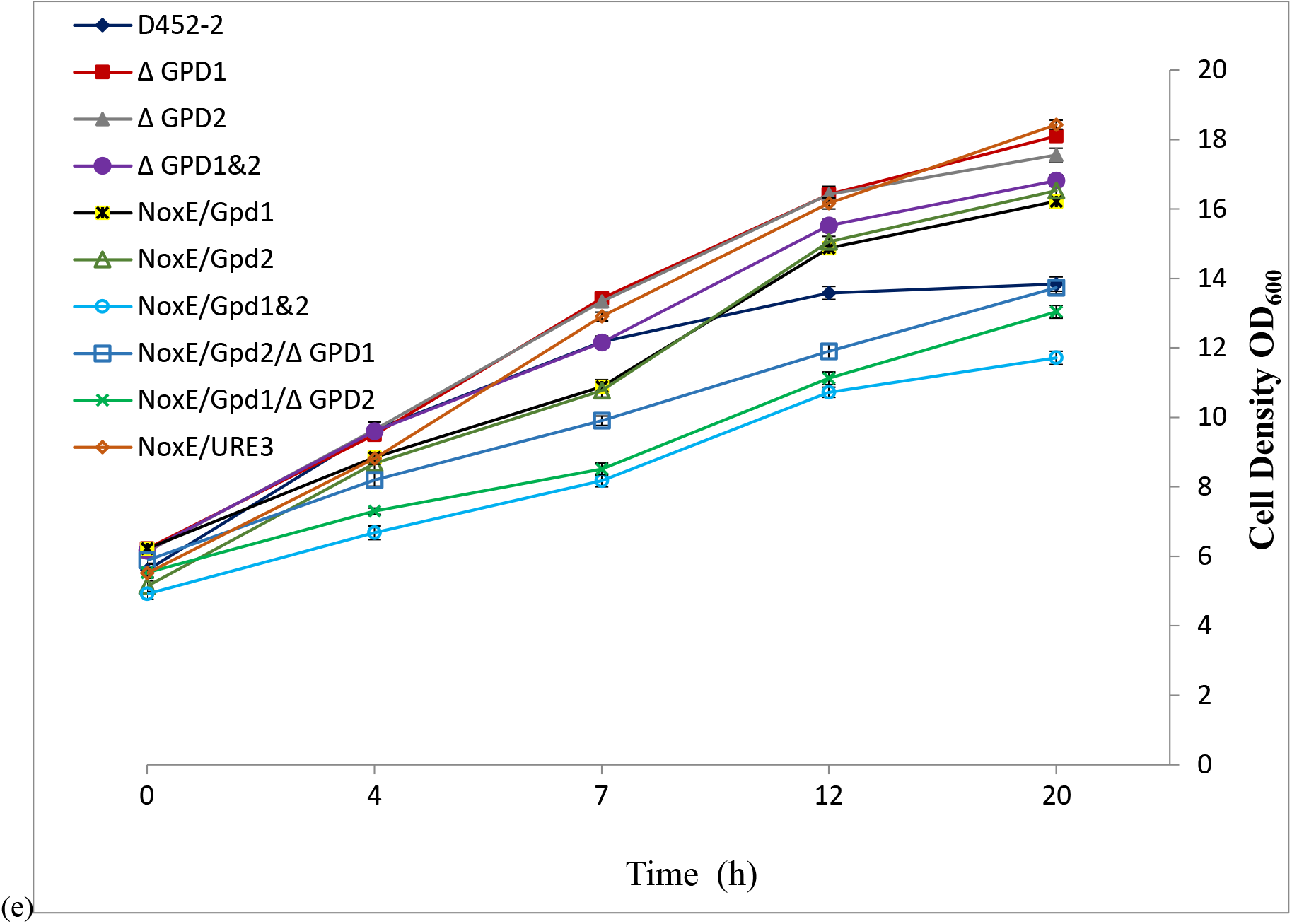
Time course for the comparative study of different recombinant *S. cerevisiae* under fermentation of 10% glucose at 30 °C and 150 rpm (1/5 liquid culture/flask volume). Data obtained from the mean of two independent experiments, (a) Glycerol production, (b) glucose consumption, (c) ethanol production, (d) acetate accumulation and (e) cell density (OD600). Data obtained from the mean of three independent experiments run at the time to decrease time-differences of sampling. Data obtained from the mean of three independent experiments run at the time to decrease time-differences of sampling. Error bars represent the standard deviation of the mean and are not visible when smaller than the symbol size. We omitted the data indicated for fermenting the only YP due to the overlapped with the horizontal axis.

**Fig. S2:**
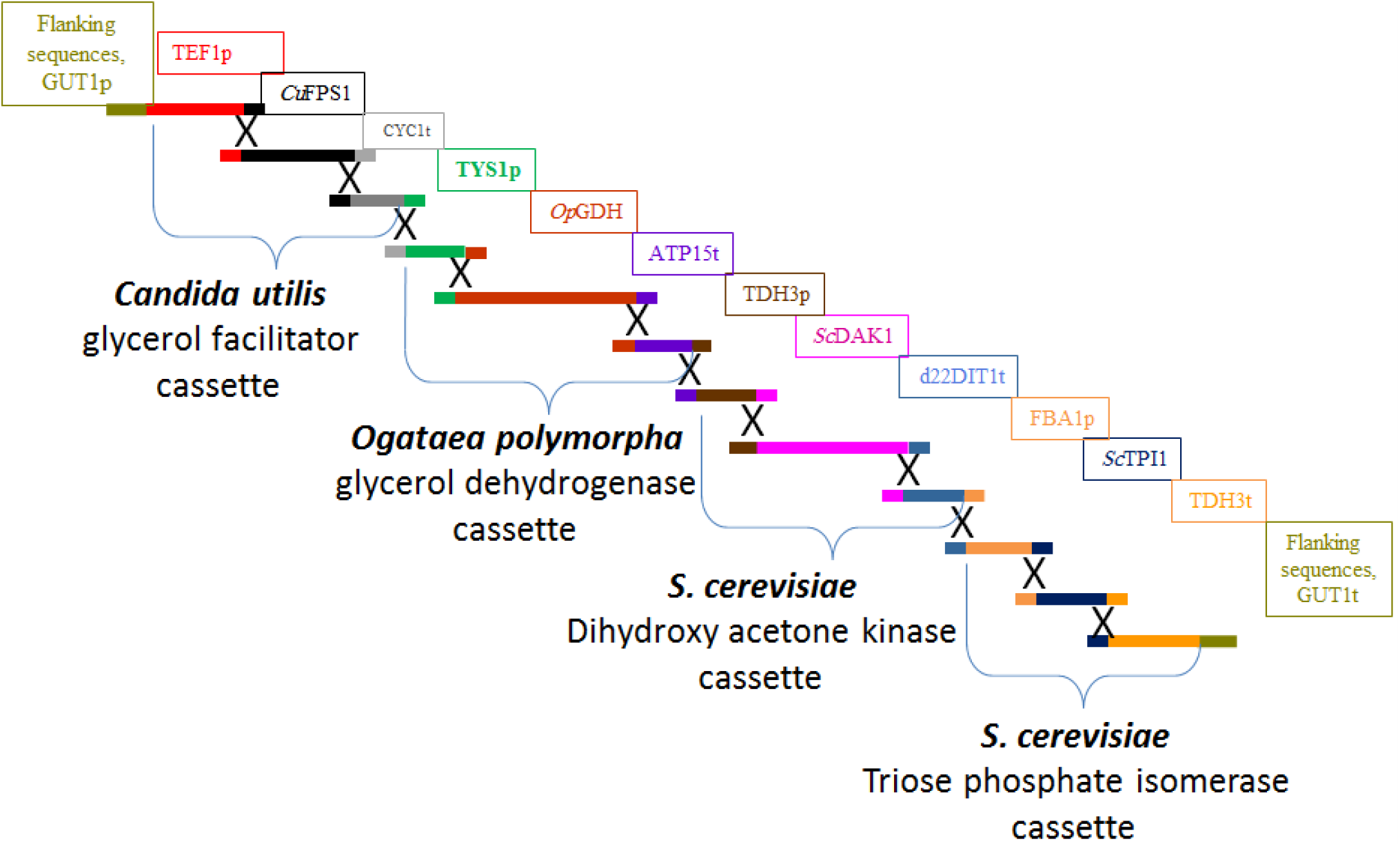
Depicted module M1 for replacing GUT1 gene by homologous recombination using CRISPR system.

**Fig. S3:**
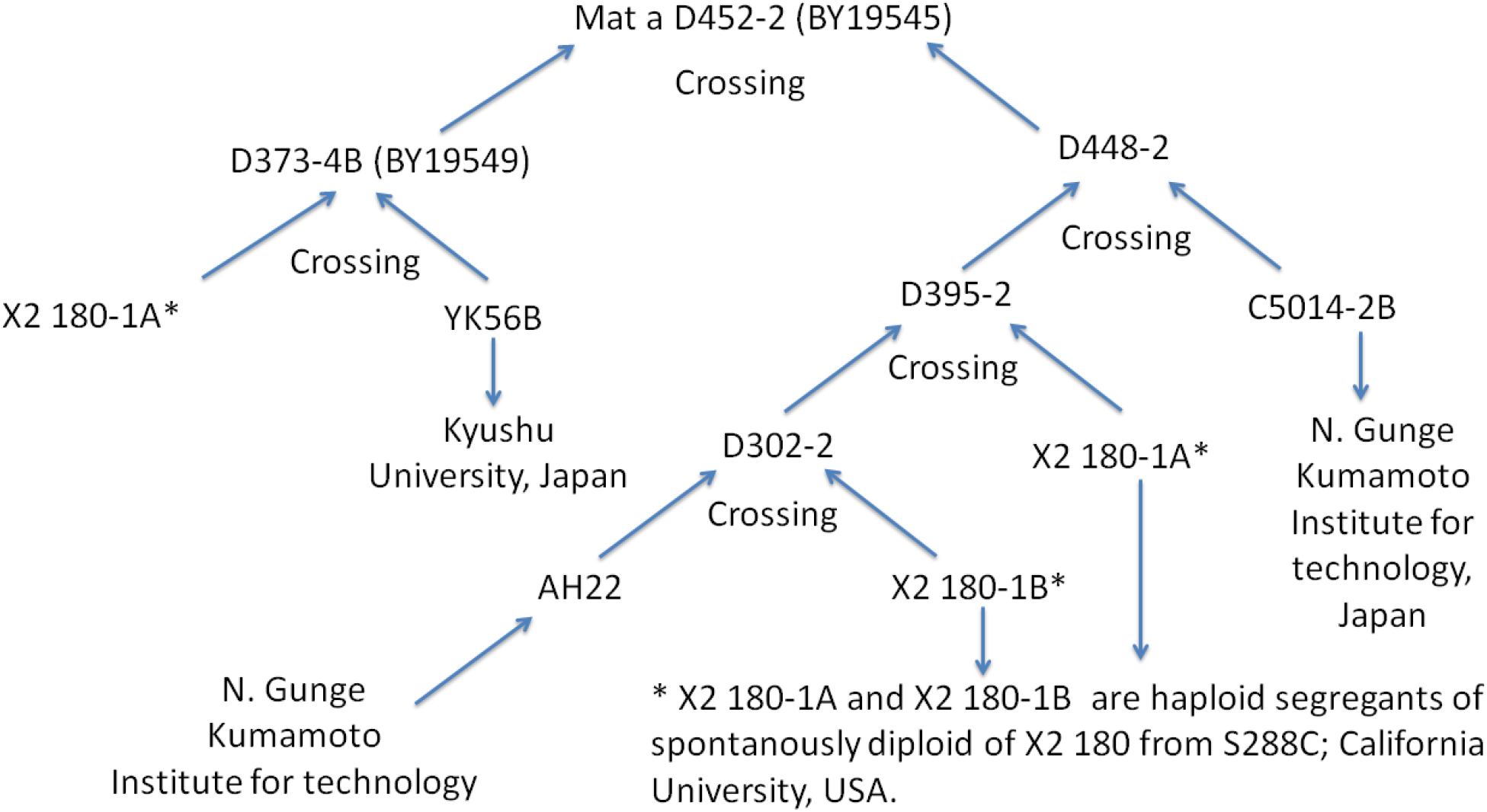
Ancestral strain with its further original^37–39^.

**Table S1.**
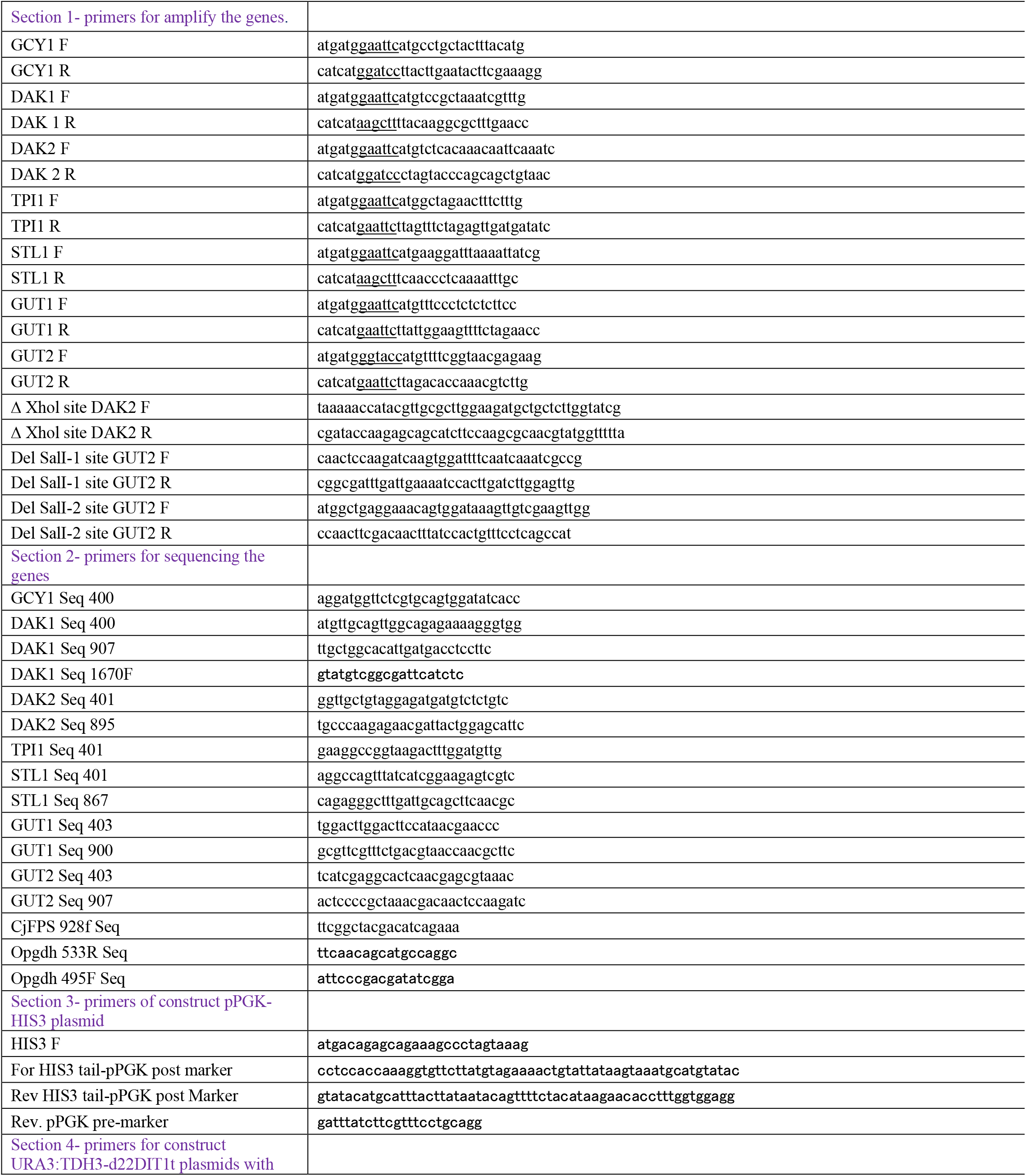
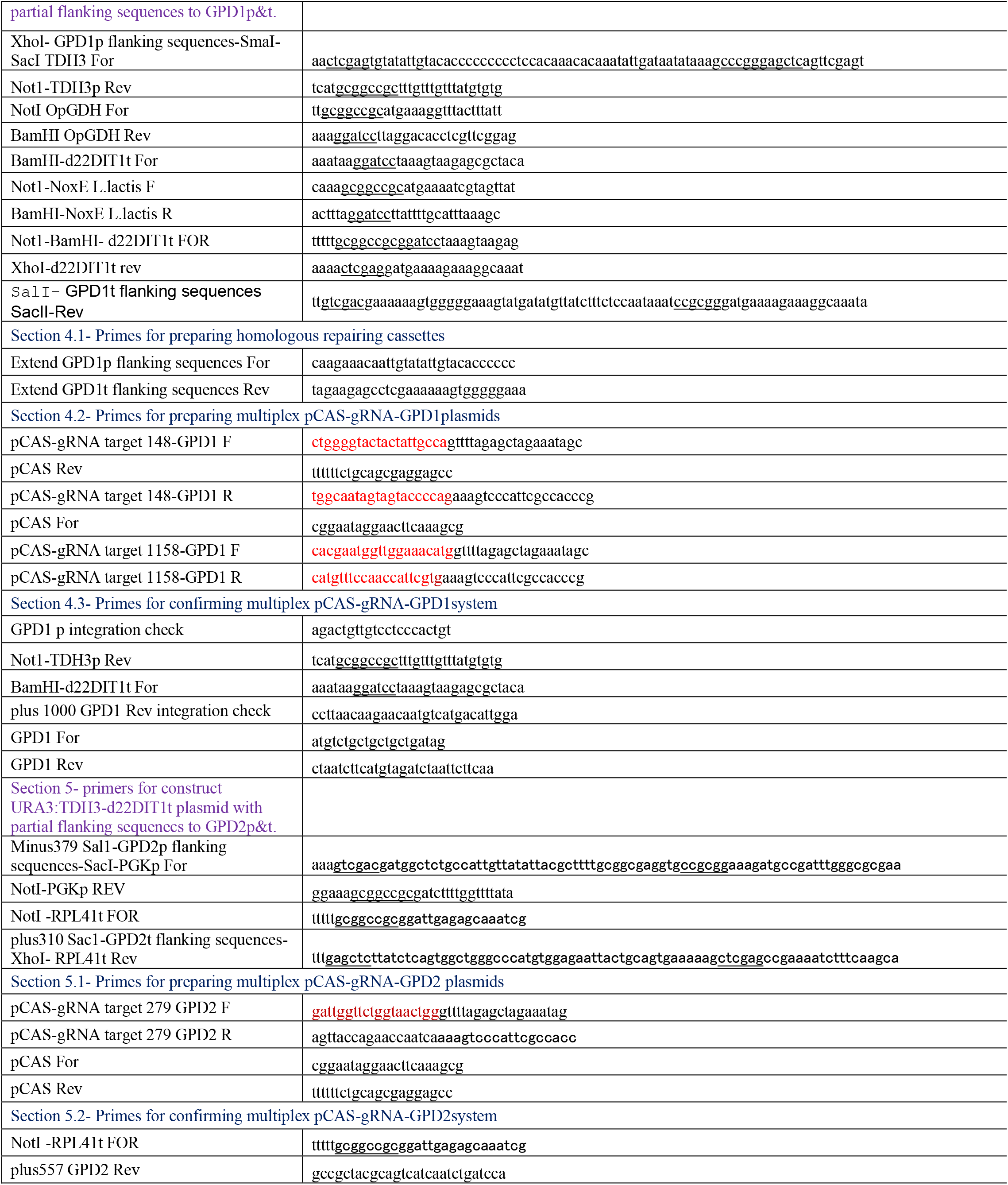
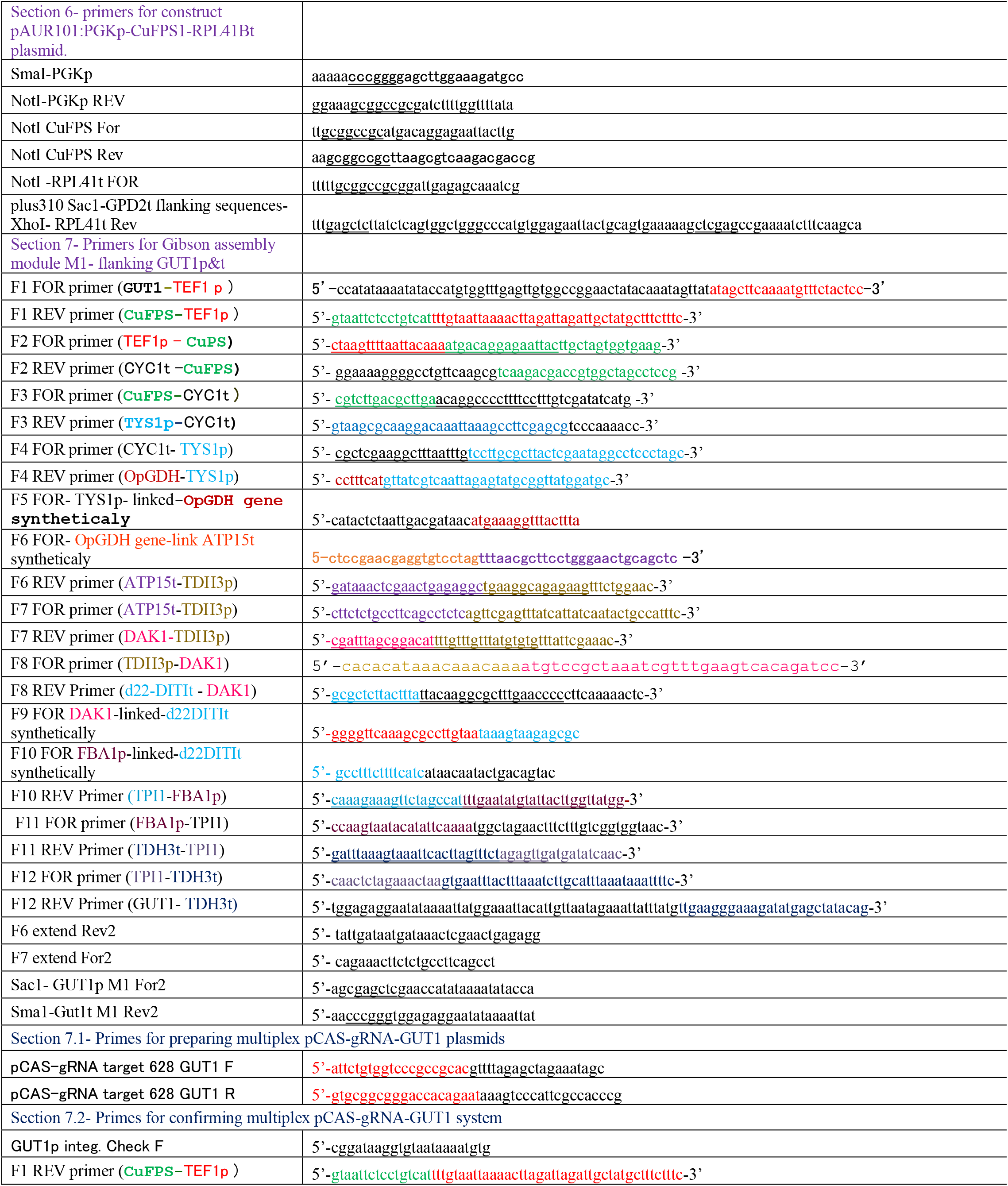
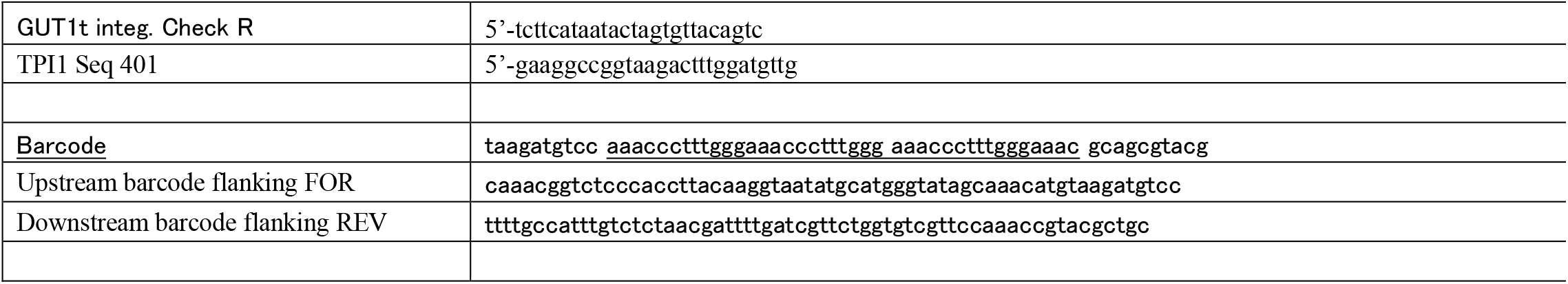
Primers used during this study.

**Table S2.**
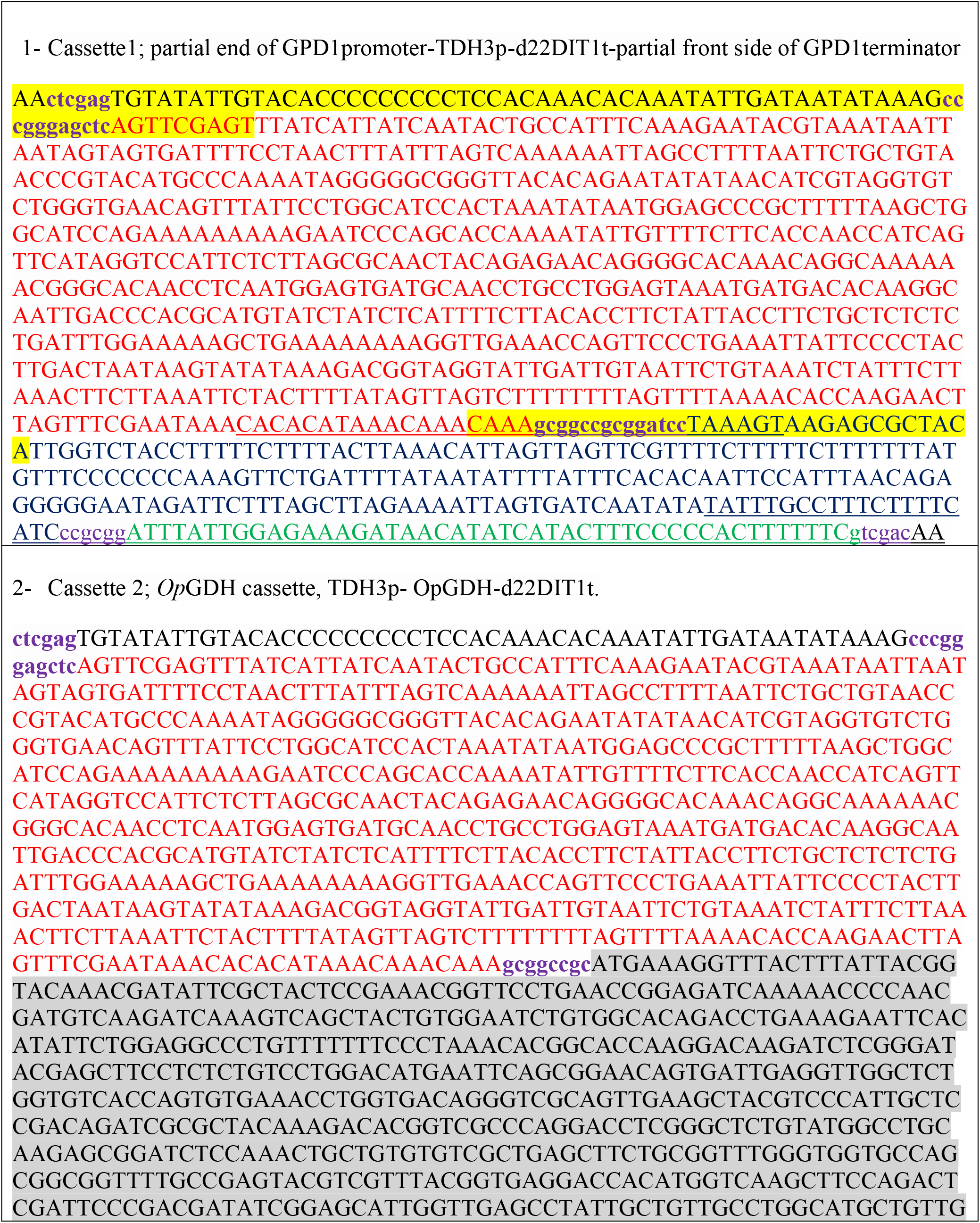
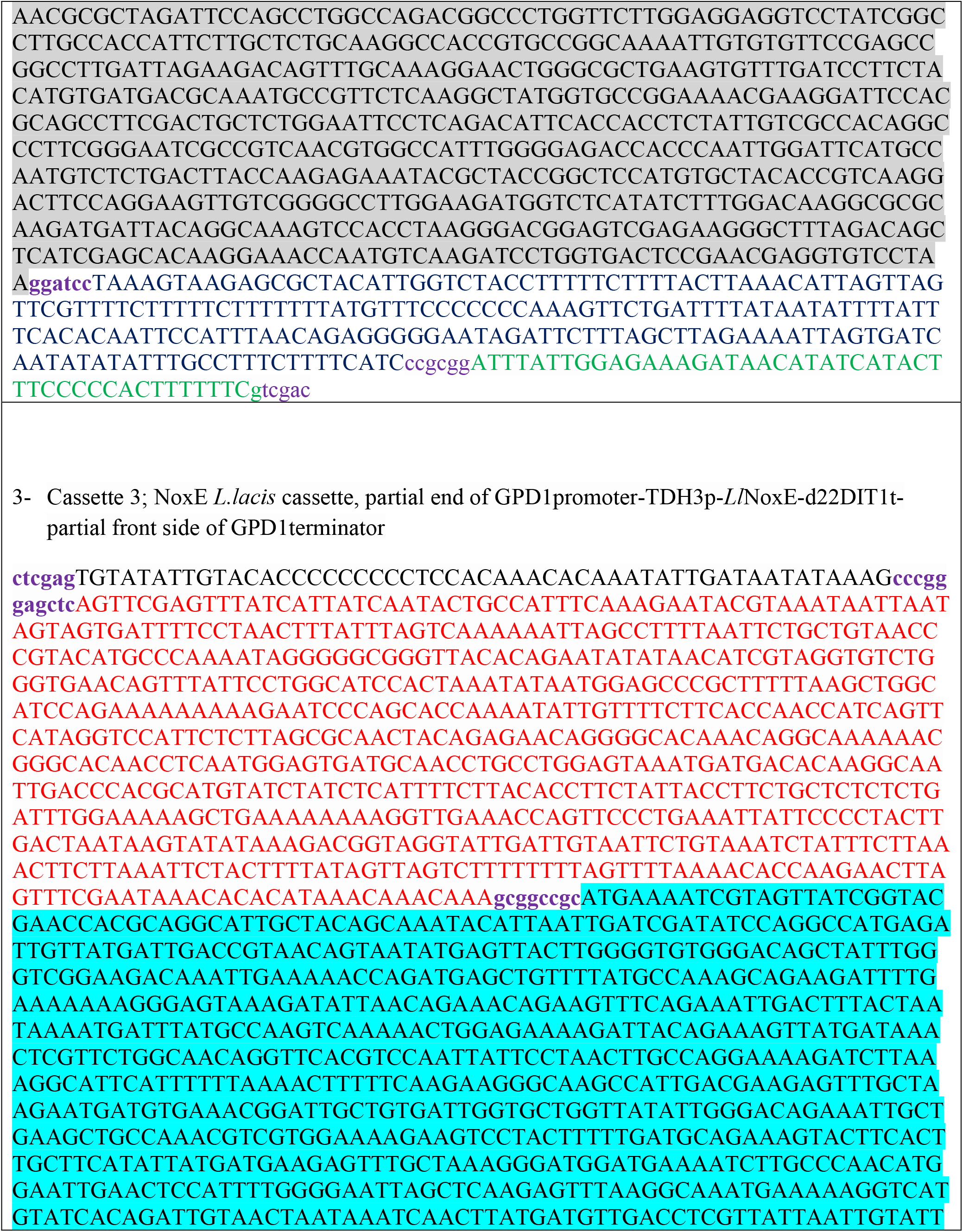
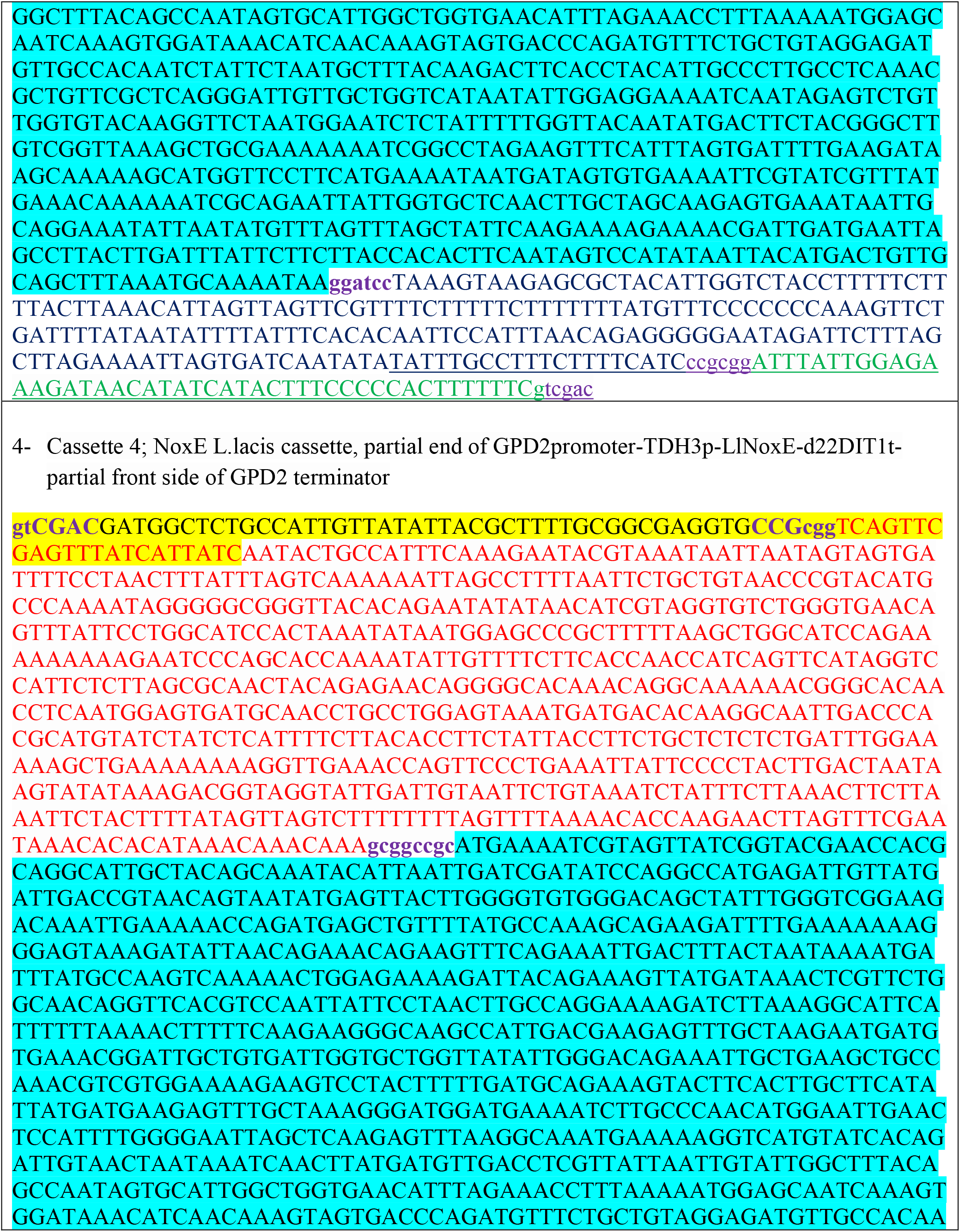
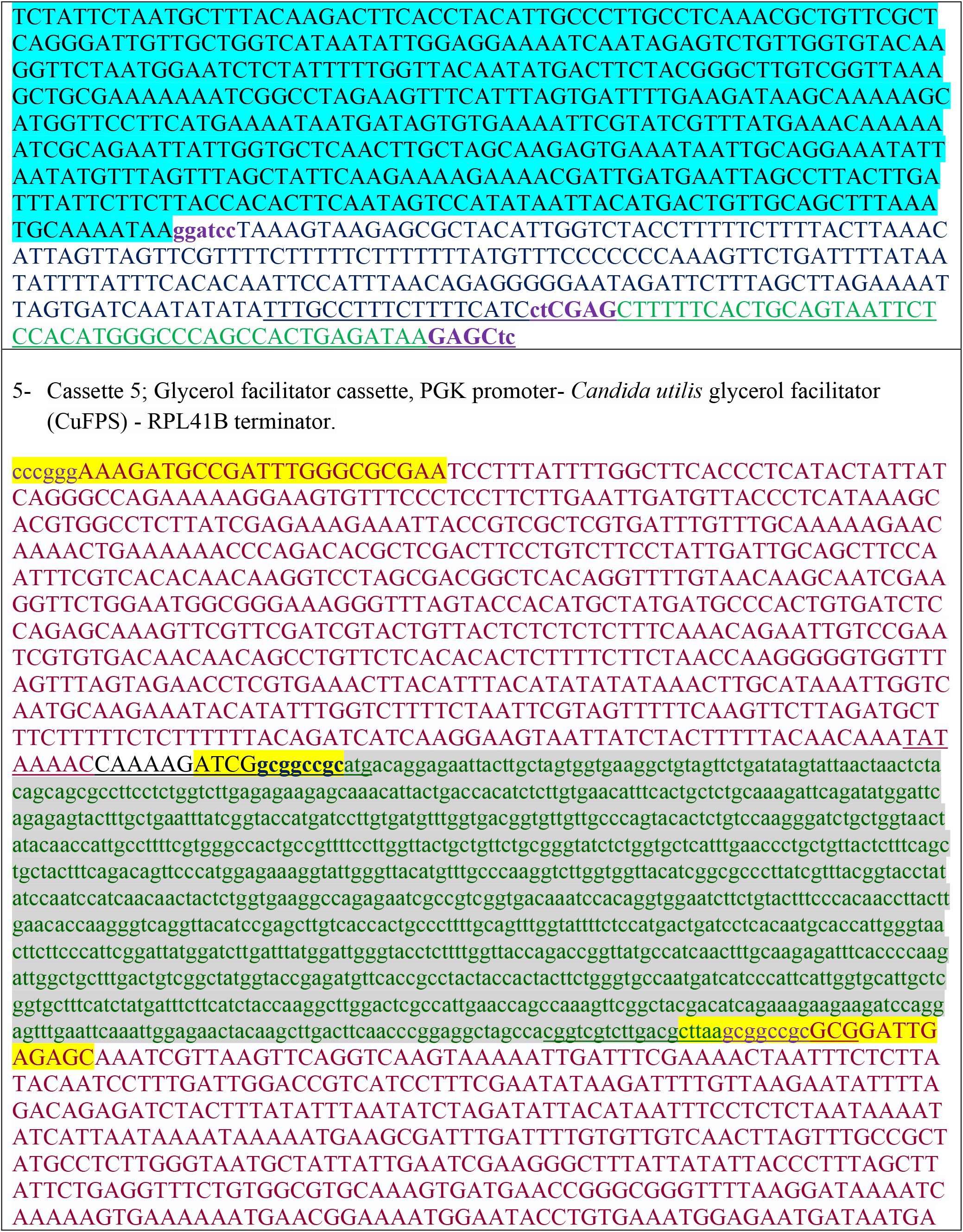
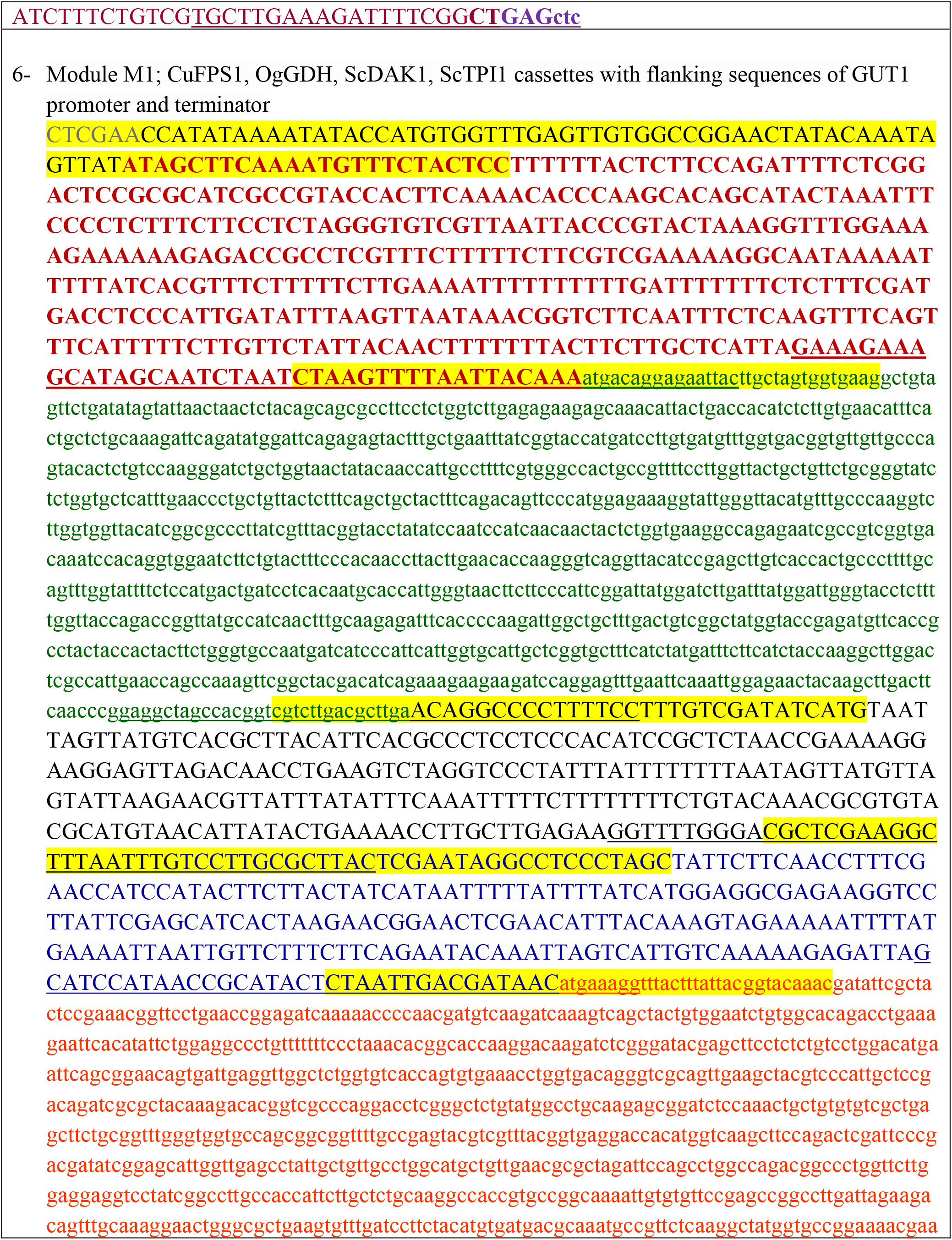
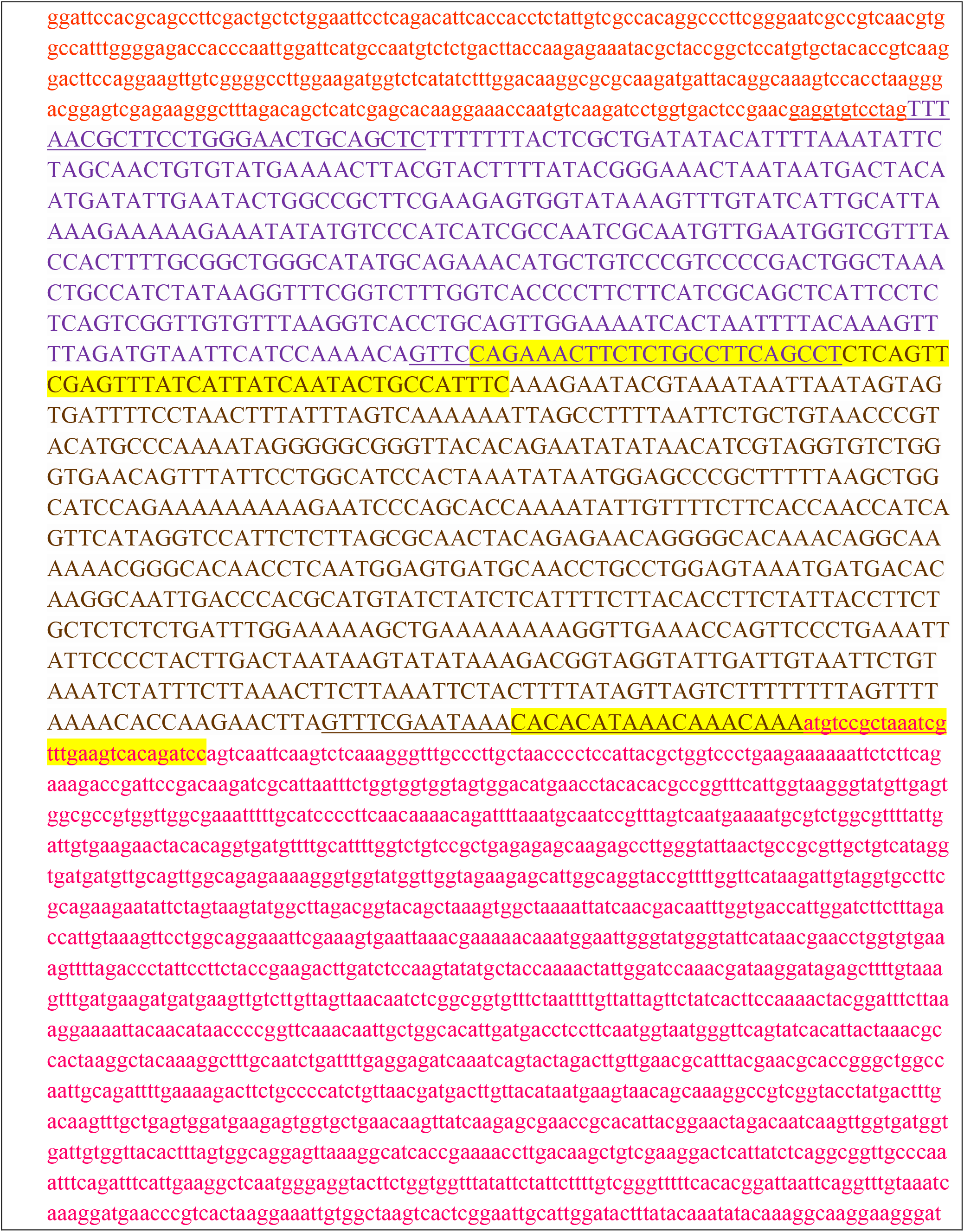
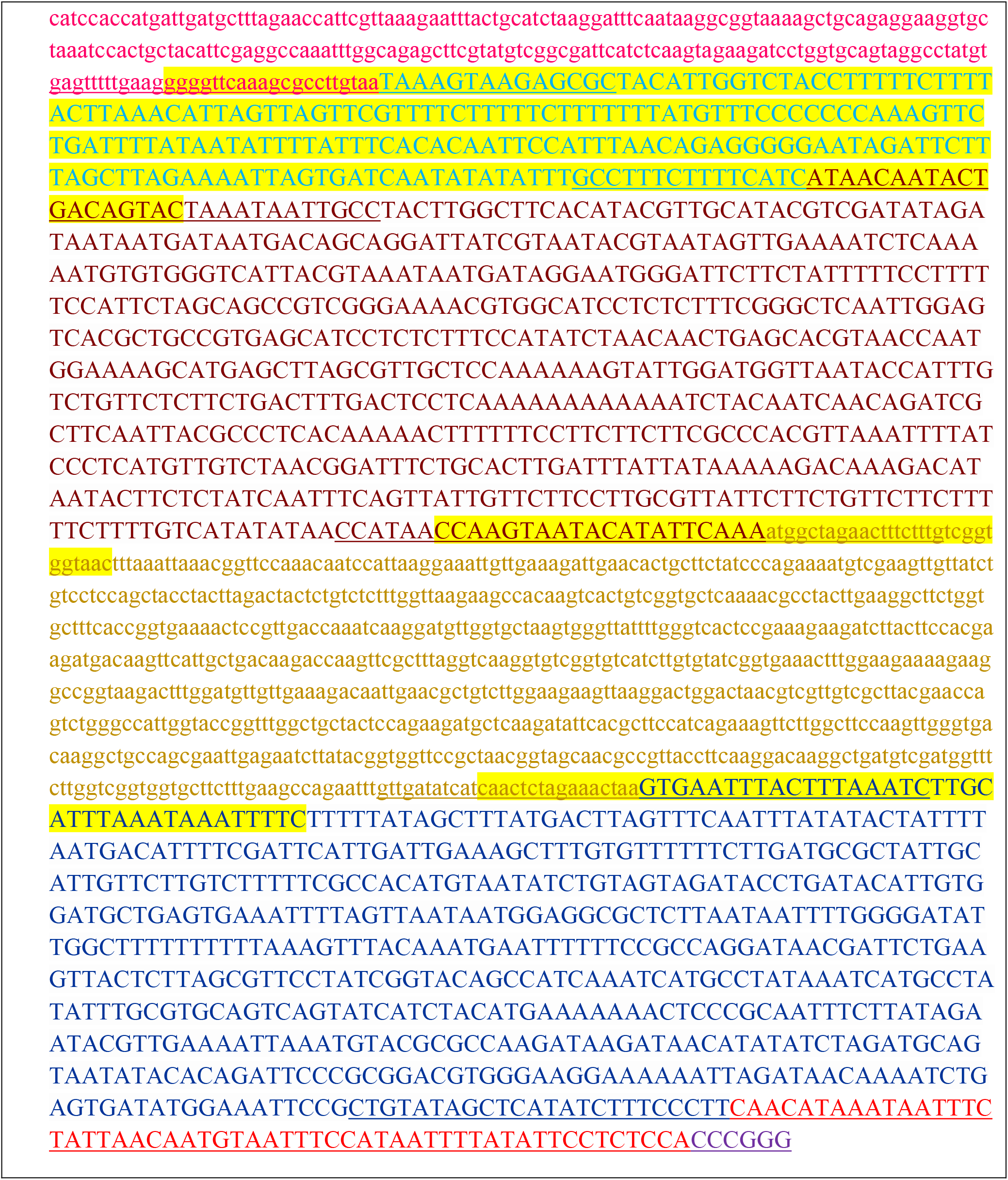
Full sequences of the integrated cassettes and module M1 with flanking sequences

